# Deep-learning models of the ascending proprioceptive pathway are subject to illusions

**DOI:** 10.1101/2025.03.15.643457

**Authors:** Adriana Perez Rotondo, Merkourios Simos, Florian David, Sebastian Pigeon, Olaf Blanke, Alexander Mathis

## Abstract

Proprioception is essential for perception and action. Like any other sense, proprioception is also subject to illusions. In this study, we model classic proprioceptive illusions in which tendon vibrations lead to biases in estimating the state of the body. We investigate these illusions with task-driven models that have been trained to infer the state of the body from distributed sensory muscle spindle inputs (primary and secondary afferents). Recent work has shown that such models exhibit representations similar to the neural code along the ascending proprioceptive pathway. Importantly, we did not train the models on illusion experiments and simulated muscle-tendon vibrations by considering their effect on primary afferents. Our results demonstrate that task-driven models are indeed susceptible to proprioceptive illusions, with the magnitude of the illusion depending on the vibration frequency. This work illustrates that primary afferents alone are sufficient to account for these classic illusions and provides a foundation for future theory-driven experiments.

## Introduction

Perception needs to turn fleeting patterns of spiking inputs into knowledge about the body and surrounding world. This challenge is formidable, as most relevant knowledge to an organism is not trivially contained in the input patterns. For instance, it is not obvious to tell from light rays hitting the retinae if the emitting object is friend or foe or if the emitting object affords to be a sittable or walkable (Gibson, 2014; Gregory, 1968). Thus, perception is a complex process turning both sensory input, as well as prior knowledge about the statistics of the world, into beliefs about the world (Dayan et al., 1995; Kawato et al., 1993; Mumford, 1994; Von Helmholtz, 1867; Yuille and Kersten, 2006). Due to the complexity of this problem, perceptual systems make systematic errors, some of which are called perceptual illusions. The nervous system is indeed subject to a number of illusions, both for specific senses and also across senses (Gregory, 1968). Apart from being fascinating phenomena, they allow to study the mechanisms and limits of conscious perception and provide essential hints about how the brain works (Geisler and Kersten, 2002; Gregory, 1968; Kanwisher et al., 2023; Weiss et al., 2002).

Proprioception is also subject to illusions (Blanke et al., 2015; Marasco and de Nooij, 2023; Prochazka, 2021; Proske and Gandevia, 2012). Vibrations of the tendon of either the biceps or triceps brachii with frequencies around 100 Hz lead to an estimated position of the elbow that is biased so as to elongate the vibrated muscle (Goodwin et al., 1972b,b; Roll and Vedel, 1982). Microneurography has confirmed that primary (type Ia) afferents are most affected by these tendon vibrations (Burke et al., 1976; Roll and Vedel, 1982; Roll et al., 1989), thus strongly suggesting that muscle spindles are the principal sensor for kinesthetic perception (Marasco and de Nooij, 2023; Proske and Gandevia, 2012). In this study, we model the ascending pathway using biomechanical models of the human arm and deep learning models of the proprioceptive pathway to investigate if “virtually” vibrating tendons will lead to biased perception.

In general, we test the hypothesis that this illusion is the necessary byproduct of perception to solve challenging inverse problems. Specifically, we ask if models of the proprioceptive pathway, optimized to localize the state of the body from muscle spindle inputs, are susceptible to biased state estimation when we apply vibrations to these inputs. We build on task-driven models of the proprioceptive pathway (Sandbrink et al., 2023; Vargas et al., 2024). These task-driven models are trained from statistics of proprioceptive inputs to infer key computational objectives. Recent work showed that models that are better at inferring the state of the body (from proprioceptive inputs) are also better at predicting neural dynamics (Vargas et al., 2024). This suggests that models of the ascending pathways, when optimized on properties of the body, give rise to powerful encoding models for explaining neural data. Do they also account for proprioceptive illusions? We find that task-driven models are indeed susceptible to vibration-induced proprioceptive illusions and that the effect of the illusion is frequency-dependent.

## Methods

To train the proprioceptive models, we used movement datasets providing naturalistic arm trajectories and muscle spindle models to generate afferent inputs. We introduce these two components and then we describe the training process and the evaluation on proprioceptive illusions.

### Movement datasets and preprocessing

Firstly, we used the proprioceptive character recognition dataset (PCR) (Sandbrink et al., 2023), which contains 2858 measured end-effector trajectories of 20 characters of the Latin alphabet sampled at 66.7 Hz. These trajectories were turned into synthetic upper-limb movements (see below).

To complement the PCR dataset with a greater diversity of (recorded) upper-limb movements, we also used the Fitness Activity Dataset with Language Instruction (FLAG3D) dataset (Tang et al., 2023), which contains full-body, 3D motion-captured pose data of 60 different fitness activities sampled at 120 Hz. We isolated the shoulder, elbow, and wrist trajectories of 7200 motion sequences, and randomly sampled four 288-frame windows from each sequence. We then slowed the samples to half speed to better match the observed velocities in the PCR dataset, filtered the keypoint sequences using a temporal Savitzky-Golay filter of order one and a window size of 25 frames, and performed an affine transformation on the keypoints to align them with the arm reference frame used in PCR.

Subsequently, to account for vibration frequencies exceeding the original sampling rates, the combined motion dataset was upsampled to 240 Hz using linear interpolation, leading to a fixed motion length of 1152 frames, or 4.8 seconds.

To summarize, the processed dataset contained 36340 arm motions (of which 6709 sourced from FLAG3D), described by one elbow and three shoulder joint angles, 3D arm keypoints extracted from the skeletal model after inverse kinematics, as well as muscle length, velocity, and acceleration properties of 25 arm muscles. Overall, 30000 motions (including 5367 FLAG3D motions) were used for training; the remaining motions were held out and used for testing.

We furthermore generated 99 arm trajectories consisting of elbow flexion and extension movements. For this Elbow flexion dataset (Ef3D) we deployed a musculoskeletal model of the human arm in OpenSim, an open-source biomechanical modeling software (Delp et al., 2007; Holzbaur et al., 2005; Saul et al., 2015). We randomized initial conditions, with shoulder elevation (39°, 99°), shoulder rotation (−6°, 54°), and elevation angle (19°, 79°) sampled independently for each configuration and held constant throughout the simulation. Elbow flexion trajectories followed sinusoidal patterns of 1 to 4 cycles, with speeds between 3°/s and 666°/s and flexion angles ranging from 45° to 130°. Muscle lengths, joint coordinates, and positions of the shoulder, elbow, and wrist were recorded over 320 time points at 66.7 Hz. To reduce noise, muscle lengths were first median-filtered, then smoothed using a Savitzky-Golay filter (window length 5, polynomial order 2), and joint positions were centered relative to the shoulder position. The dataset comprising 20 motions was then upsampled to 240 Hz using linear interpolation to ensure compatibility with the combined motion dataset.

To test muscle-tendon vibrations we generated 100 arm trajectories consisting of static poses in different arm configurations. For this Elbow static dataset (Es3D), we followed the same approach as for Ef3D to generate the initial arm configuration but maintained the arm static for the totality of the trial.

### Computing joint angles and muscle length trajectories

We followed the method presented in Sandbrink et al. (2023) to transform elbow-wrist keypoint sequences into length trajectories of 25 arm muscles. Firstly, we used a skeletal model of the arm and performed inverse kinematics to extract the elbow flexion and shoulder elevation, adduction, and rotation joint angles from the 3D shoulder, elbow, and wrist positions. Following that, we deployed a musculoskeletal model of the human arm by Holzbaur et al. (2005); Saul et al. (2015) in OpenSim, an open-source biomechanical modeling software (Delp et al., 2007), and extracted the length properties at equilibrium of 25 arm muscles corresponding to the extracted joint angle sequences. This process was performed for each time frame across all movement windows for both the PCR and the extracted FLAG3D motions. For PCR, we used the provided data (Table 1) while for FLAG3D, Es3D and Ef3D we adapted the code to generate the muscle state trajectories (Table 1).

**Table 1.** Summary of software and datasets used and generated in this study.

| Resource | Reference | Link |
| --- | --- | --- |
| **Datasets** |  |  |
| Synthetic muscle spindle datasets and models | This paper | https://github.com/amathislab/ProprioceptiveIllusions |
| PCR dataset | Sandbrink et al. (2023) | https://zenodo.org/records/14544688 |
| FLAG3D dataset | Tang et al. (2023) | https://andytang15.github.io/FLAG3D |
| **Software and algorithms** |  |  |
| Code | This paper | https://github.com/amathislab/ProprioceptiveIllusions |
| TensorFlow variant | Sandbrink et al. (2023) | https://github.com/amathislab/DeepDraw |
| PyTorch | Paszke et al. (2019) | https://pytorch.org |

Finally, we computed the first and second moments of the extracted muscle length sequences (velocity and acceleration) smoothed using a Savitzky-Golay filter of order one and window size of 31 frames, which we used as an approximation of the dynamic properties of the muscles. All smoothing steps were performed to remove movement artifacts and address potential discontinuities arising at various processing steps. Movement sequences that contained such artifacts were removed, thus ensuring that all data was accurate and within reasonable range.

To ensure uniformity across the 25 muscles with varying lengths, we normalized muscle lengths relative to their optimal fiber length *L*_0_ provided by the musculoskeletal model to obtain the normalized muscle length *l*(*t*) = *L*(*t*)*/L*_0_ where *L*(*t*) is the fiber length provided by the simulation environment.

### Modeling muscle spindles

Following Dimitriou and Edin (2008); Prochazka and Gorassini (1998), we assumed a transfer function that is either linear or includes a power term in the velocity component. In our previous work on modeling the proprioceptive pathway (Sandbrink et al., 2023; Vargas et al., 2024), we also based our spindle models on those works. However, we had made the simplifying argument that one can provide the state of the muscles to the deep learning models via two input channels^1^ per muscle: muscle length and velocity. Due to the large number of afferents, the system could effectively demix muscle length and velocity, which is why we directly provided those. In an earlier version of Sandbrink et al. we used the model by Prochazka and Gorassini (1998) and observed the emergence of similar tuning curves (see v1 on Biorxiv).

Here we deviated from this practice to allow the modeling of responses to muscle-tendon vibrations. These responses need to be mediated through afferents that combine all those signals. The muscle spindle responses to normalized muscle length *l*, velocity *v*, and acceleration *a* were modeled using a linear transfer function with a velocity power term. The firing rate *f* for primary (type Ia) and secondary (type II) afferents was expressed as:

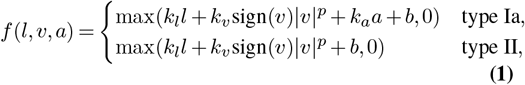

where *k*_*l*_, *k*_*v*_, *k*_*a*_, *b*, and *p* are scalar parameters. For type II afferents, the acceleration term *a* was excluded due to their reduced sensitivity to velocity changes (Proske and Gandevia, 2012). The velocity power coefficient *p* can be set to either 0.6 or 1, reflecting values found in the literature (Dimitriou and Edin, 2008; Prochazka and Gorassini, 1998). We note that in this article we use *p* = 1 throughout.

### Optimizing spindle model parameters

To ensure biologically realistic behavior, the spindle model was constrained in two ways. First, the firing rates were required to remain within the range [0, *f*_max_] for all possible arm configurations in the movement dataset. Second, each afferent was constrained to exhibit a certain level of lifetime sparseness quantified by *λ*_*s*_, i.e., the proportion of time points where the firing rate was nonzero (Willmore and Tolhurst, 2001).

The proprioceptive models were trained with afferent inputs generated by optimizing spindle model parameters. For each muscle, a set of *N* afferents (we used 5 from each type) was modeled. The maximum firing rate (*f*_max_) for each afferent was drawn from a probability distribution based on findings by Roll et al. (1989). For type Ia afferents, *f*_max_ followed a uniform distribution between 50 and 180 Hz. For type II afferents, the maximum firing rate was sampled from a uniform distribution between 20 and 50 Hz. Additionally, a sparseness level *λ*_*s*_ was sampled from a uniform distribution between 70 and 100%.

Once these parameters were drawn, the spindle model parameters *θ*_*s*_ = {*k*_*l*_, *k*_*v*_, *k*_*a*_, *b*} were optimized using a subset of 2000 movements from the FLAG3D dataset, which we concatenated. The optimization was performed by minimizing a cost function 𝒞 (*θ*_*s*_), which penalized the deviation of the predicted firing rate from the maximum firing rate *f*_max_ and the sparseness of the afferent firing rates. The cost function is given by:

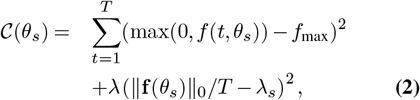

where *f* (*t, θ*_*s*_) represents the predicted firing rate at time *t* for the model with parameters *θ*_*s*_, and **f** (*θ*_*s*_) is the vector of firing rates across all time points. The sparseness function ∥.∥ _0_*/T* computes the fraction of time points where the firing rate of the muscle spindle is nonzero. A multiplicative constant *λ* was chosen empirically to aid convergence of the optimization.

After optimization, the optimal parameters *θ*_*s*_ for each afferent were saved. Due to the non-convexity of the objective function, the solutions are not unique. We could thus generate independent parameters across afferents. In addition, we recorded the actual sparseness and maximum firing rate of each afferent across the movement dataset. Overall, we computed the optimal parameters for 30 independent type Ia afferents and for 30 independent type II afferents. We then defined five coefficient sets by randomly sampling five type Ia and five type II afferents for each muscle.

### Training proprioceptive models

We considered models that predict the postural state of the body from the distributed inputs of muscle spindles. Those were among the best for predicting neural data in the somatosensory brain stem and cortex (Vargas et al., 2024). Furthermore, the architecture search from Sandbrink et al. (2023) showed that spatiotemporal models with sufficiently many layers could accurately estimate the state of the arm (location). There are three key differences in the models trained here. First, to account for vibrations, the sampling rate for both inputs and outputs was increased. Second, instead of using muscle length and velocity, the model received spindle afferent signals as input.

Third, the model predicted not only the location of the end-effector from spindle afferents, but also joint angles (one for the elbow and three for the shoulder). We note that the coordinate framework had little impact on the neural predictability (Vargas et al., 2024). We modified the original TensorFlow implementation from Sandbrink et al. (2023) to PyTorch (Paszke et al., 2019), incorporating the changes described here.

We shaped the afferent signal by including 2*N* time-series of type Ia and type II firing rates (*N* for each type), for each of the 25 arm muscles. These time-series translate to a total input size of *B ×* 2*N ×* 25 × *T* input with *B* denoting the batch size and *T* = 1152 denoting the sequence length in frames. The neural network consists of four convolutional layers, each with a 7×7 kernel, stride of 1, padding of 3, applied along the spatiotemporal dimensions of the input data. The spindle afferent channels change size from 2*N* to 8, 8, 32, and 64. The output of each convolutional layer is passed through a ReLU activation function. The output of the last convolutional layer is flattened in the muscles and afferent dimensions. A final linear layer outputs the predicted end-effector position in XYZ along with the four joint angles of the arm, processing each time point individually.

Similarly to Sandbrink et al. (2023), the spatiotemporal model was trained using batch gradient descent with an Adam optimizer with an initial learning rate of 0.0005 and decay parameters of 0.9 and 0.999 to minimize the mean squared error between predicted and true trajectories of the predicted end-effector and joint trajectories. We set the batch size to 128 and the number of epochs to 1000. Early stopping was implemented based on loss in the validation set. The learning rate was divided by four if the validation loss did not improve over four epochs. Training was terminated if performance did not improve over two early stopping cycles and a minimum of 40 epochs had already elapsed. To facilitate training, we computed the mean and standard deviation of the training dataset to normalize the inputs and outputs of the neural network. These values were used during inference to scale and un-scale the inputs and outputs, respectively.

Once the afferent parameters were optimized, the proprioceptive model was trained using the entire training set (without vibrations) and monitored with the validation set. Performance was evaluated on a separate test set after training. The datasets were split similarly to Sandbrink et al. (2023), where samples were randomly assigned to training, validation, and testing sets with a 72-8-20 ratio, respectively. The accuracy of the model was defined as the decoding error, expressed as the mean Euclidean distance between the predicted and true wrist coordinates. Additionally, the elbow angle accuracy was computed as the root mean squared error between the predicted and true elbow angles.

### Modeling proprioceptive illusions

#### Effect of tendon vibrations on proprioceptive inputs

The effect of tendon vibrations on proprioceptive inputs was modeled according to microneurography studies (Burke et al., 1976; Roll and Vedel, 1982; Roll et al., 1989). First, the arm position was set to an initial configuration, and the corresponding spindle responses **s** for this configuration, assuming no vibrations, were generated. Second, a vibration frequency *ν* within the range [0, 190] Hz was applied to selected muscles (typically either both biceps or the three triceps heads). For afferents of type Ia (and type II when vibrated), the perturbed firing rate was computed using the following function:

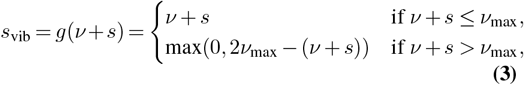

where *ν*_max_ represents the maximum harmonic frequency of the spindle and *s* the spindle response to the state of the arm without vibration. This expression is based on prior findings by Roll et al. (1989) showing that the firing rate of afferents exhibits a nonlinear relationship with the vibration frequency, with their firing rate being locked one-to-one to the vibration frequency until an afferent-specific maximum vibration frequency. For each afferent, we initially assumed the maximum vibration frequency corresponds to the maximum firing rate *ν*_*max*_ = *f*_*max*_ used to find the spindle parameters, as explained in the subsection “Optimizing spindle model parameters.” We later explored the effect of the maximum vibration frequency on the illusion by setting *ν*_*max*_ to a specific value for all afferents or by drawing it from a uniform distribution (Figure 5).

Unless otherwise specified, type II afferents were assumed to be unaffected by tendon vibrations, consistent with experimental observations that secondary afferents are less sensitive to vibrations than primary ones, especially for low-amplitude vibrations (Burke et al., 1976; Roll and Vedel, 1982; Roll et al., 1989). Their firing rates remained unchanged under vibratory conditions and were expressed as:

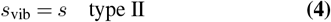

where *s* is the response of the afferent without vibration. When we applied vibrations to type II afferents (Figure 6), we used the same transfer function as for type Ia Eq. (3) using the afferent maximum firing rate as the maximum harmonic frequency *ν*_*max*_ = *f*_*max*_.

To quantify the effect of tendon vibrations on elbow angle estimation, we used the Es3D dataset, comprising 100 static arm configurations. Tendon vibrations were applied to the muscles between *t*_*s*_ ≙ 200 steps = 0.83*s* and *t*_*e*_ = 900 steps ≙ 3. 75 *s*, to include for each trial a period before and after vibration. For each trial, we calculated the perceived angle difference (or “illusion” angle) as the change in predicted elbow angle between the pre- and during-vibration periods by comparing the mean angle over 10 steps immediately before the vibration (*t*_*s*_ *−* 40 steps and *t*_*s*_ *−* 30 steps) to the mean angle over 10 steps during the vibration (*t*_*s*_ + 100 steps to *t*_*s*_ + 110 steps).

#### Vibration of other muscles

To evaluate how the elbow angle illusion varies when targeting different muscles, we applied 100 Hz vibrations to the type Ia afferents of each of the 25 modeled muscles individually (Table 2). We repeated this experiment for all coefficient seeds and all trained models (5 coefficient seeds, 20 trained models in total), recording the perceived elbow angle difference in each case. To interpret the variations in the magnitude of the illusion across muscles, we characterized each modeled muscle based on its functional relationship to the elbow joint angle (Figure 7). Additionally, we computed the Pearson correlation between the afferent firing rates of each muscle and the elbow angle throughout the training dataset (10 spindles per muscle, 25 muscles, 5 spindle coefficient sets, 30000 × 1152 timepoints).

**Table 2.**
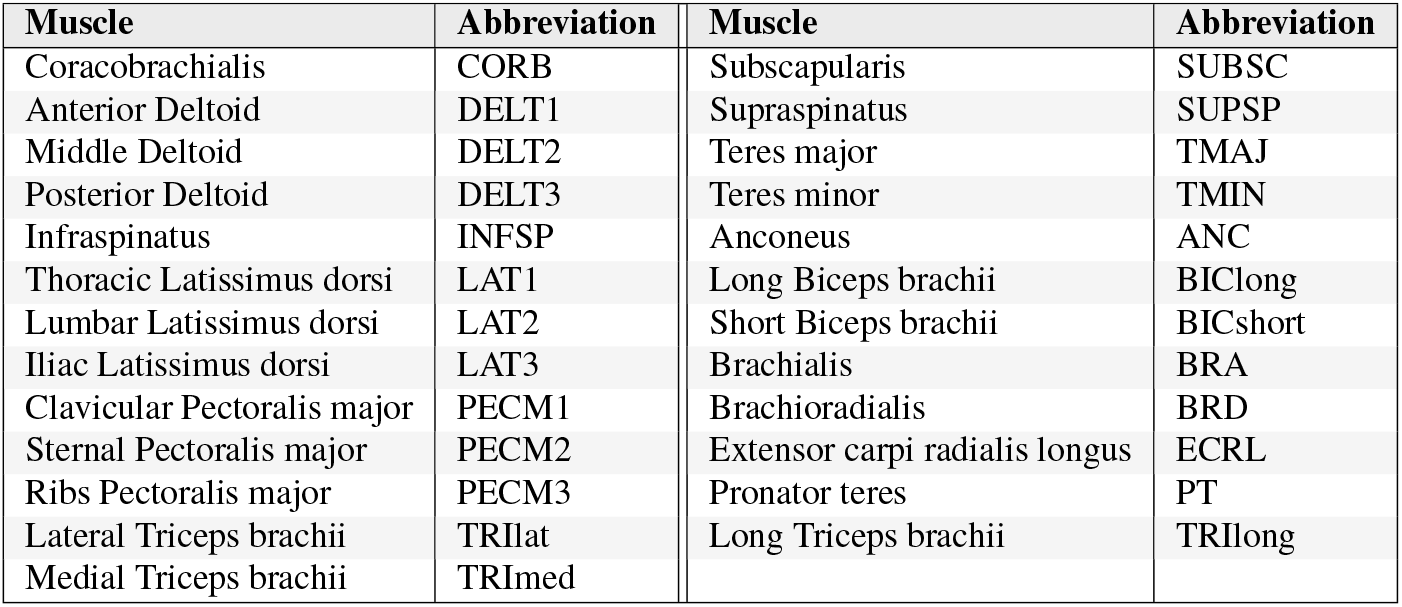
List of arm muscles used in the model and their abbreviations.

| Muscle | Abbreviation | Muscle | Abbreviation |
| --- | --- | --- | --- |
| Coracobrachialis | CORB | Subscapularis | SUBSC |
| Anterior Deltoid | DELT1 | Supraspinatus | SUPSP |
| Middle Deltoid | DELT2 | Teres major | TMAJ |
| Posterior Deltoid | DELT3 | Teres minor | TMIN |
| Infraspinatus | INFSP | Anconeus | ANC |
| Thoracic Latissimus dorsi | LAT1 | Long Biceps brachii | BIClong |
| Lumbar Latissimus dorsi | LAT2 | Short Biceps brachii | BICshort |
| Iliac Latissimus dorsi | LAT3 | Brachialis | BRA |
| Clavicular Pectoralis major | PECM1 | Brachioradialis | BRD |
| Sternal Pectoralis major | PECM2 | Extensor carpi radialis longus | ECRL |
| Ribs Pectoralis major | PECM3 | Pronator teres | PT |
| Lateral Triceps brachii | TRIlat | Long Triceps brachii | TRIlong |
| Medial Triceps brachii | TRImed |  |  |

All the code is available (Table 1).

## Results

We sought to model perceptual illusions related to tendon vibrations (Eklund, 1972; Goodwin et al., 1972b,b; Roll and Vedel, 1982). For instance, during a stationary posture when the triceps tendon were vibrated at approximately 100 Hz, subjects perceived their arm moving at the elbow joint, creating a systematic mismatch between the actual and perceived arm position Roll and Vedel (1982). The illusion biases perception toward a position where the vibrated muscle appears elongated (Figure 1A). The illusion is quantified by having subjects track their vibrated arm’s position with their unperturbed arm, thereby measuring the discrepancy in perceived elbow angle. As most illusions are described for arm muscles, we focused on modeling the ascending proprioceptive pathway of the human arm comprising 25 muscles (Holzbaur et al., 2005; Saul et al., 2015) and assessed illusion effects by measuring how vibration altered the predicted elbow angle.

**Figure 1.**
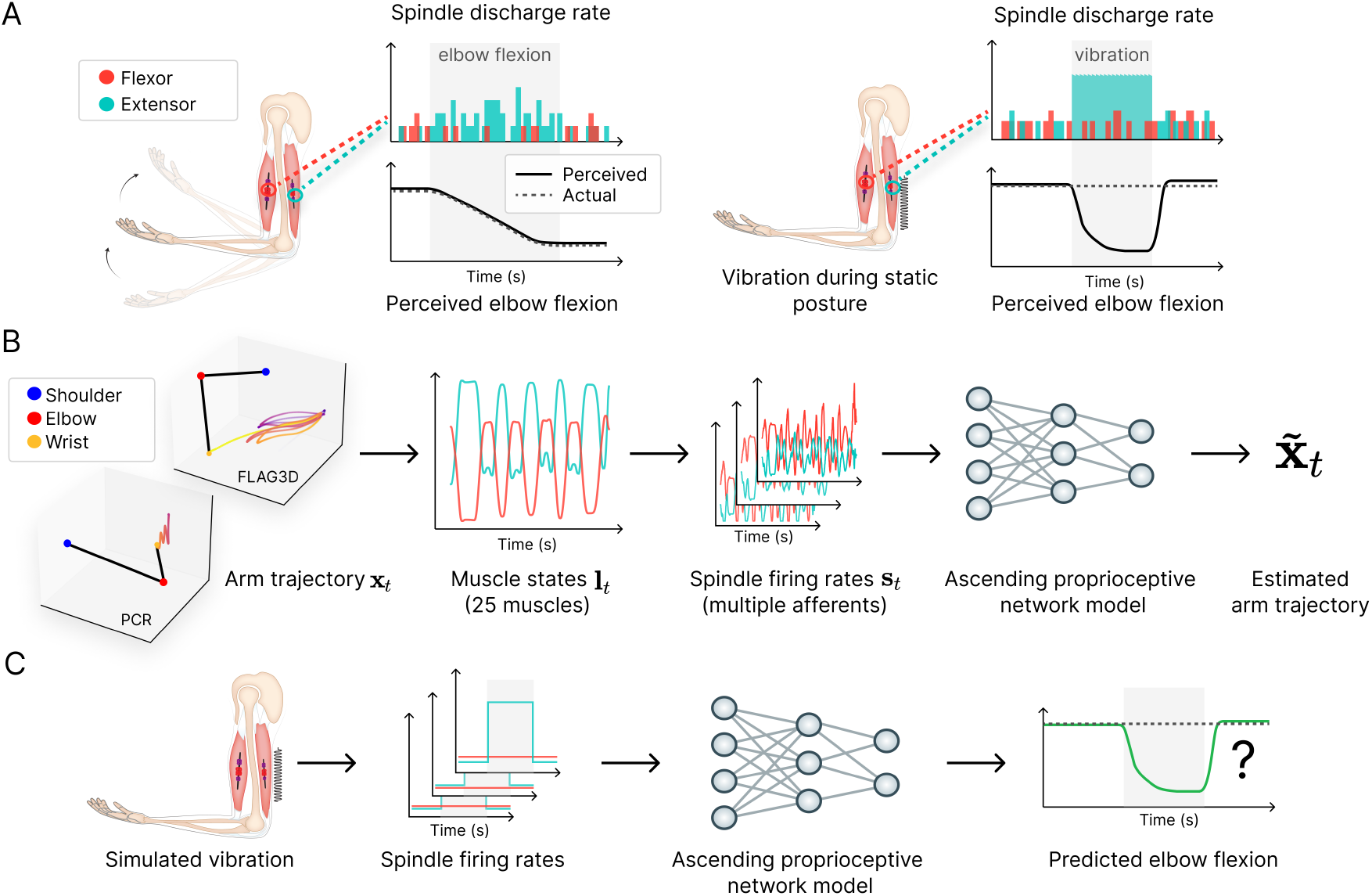
**A** Afferent signals from muscle spindles facilitate the accurate perception of movement. Eliciting an artificial muscle spindle response through muscle-tendon vibration can lead to an illusory perception of movement. Arm graphic adapted with permission from Marasco and de Nooij (2023). **B** Arm trajectories correspond to changes in muscle state, which drive muscle spindle response. We trained models of the ascending proprioceptive pathway to estimate the arm trajectories from these spindle afferents. We also trained models with different movement (and thus spindle) statistics. We show one example from the PCR and one example from the FLAG3D dataset. **C** We tested the (frozen) trained model on the vibration experiment. Does the model predict an illusory arm displacement, similar to humans? Importantly, we did not train the models with any illusion data but rather tested whether illusions are a consequence of task training.

### Mathematical motivation

The (simplified) function of the ascending proprioceptive pathway can be reasonably distilled into the following mathematical formulation (Figure 1B). Let **l**_*t*_ denote a sequence of muscle states across time, that is, the product of the upper limb being passively displaced along a trajectory **x**_*t*_. These muscle states drive specific muscle spindle responses **s**(**l**_*t*_). We can thus formulate the ascending pathway as the function 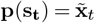, meaning that the proprioceptive neural circuit infers the state of the body from the distributed muscle spindle information. We modeled **p** by defining a neural network *M*_*θ*_, and setting the parameters *θ* according to the optimization problem

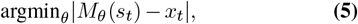

under a wide range of arm movements (movement statistics). This model of the proprioceptive pathway *M*_*θ*_ builds on the framework introduced by Sandbrink et al. (2023) (see Methods).

Our core question was: How does the model behave under muscle spindle vibration (Figure 1C)? In particular, does the model recapitulate the systematic bias observed in human psychophysics studies? Furthermore, we investigated whether the model’s behavior is robust to reasonable changes in the parameters of muscle spindles. We emphasize that our model makes many simplifications (see Discussion), and in particular only focuses on the ascending pathway. Next, we introduce the different components in more detail.

### Statistics of arm movement

As we just outlined, to constrain models of the ascending proprioceptive pathway we need statistics of arm movement. We considered two datasets. Firstly, building on Sandbrink et al. (2023) we used the PCR dataset (see Methods). Secondly, we enriched the training dataset by incorporating a more diverse array of movements, including data from the FLAG3D dataset (Tang et al., 2023), which features fitness exercises spanning a wide physiological range. Indeed, the FLAG3D dataset covers a larger portion of the reachable space, which translates to a wider, but still physiologically plausible range of muscle states (Figure 2A). Using models of muscle spindles, we generated spindle firing rates, which serve as the input to *M*_*θ*_.

**Figure 2.**
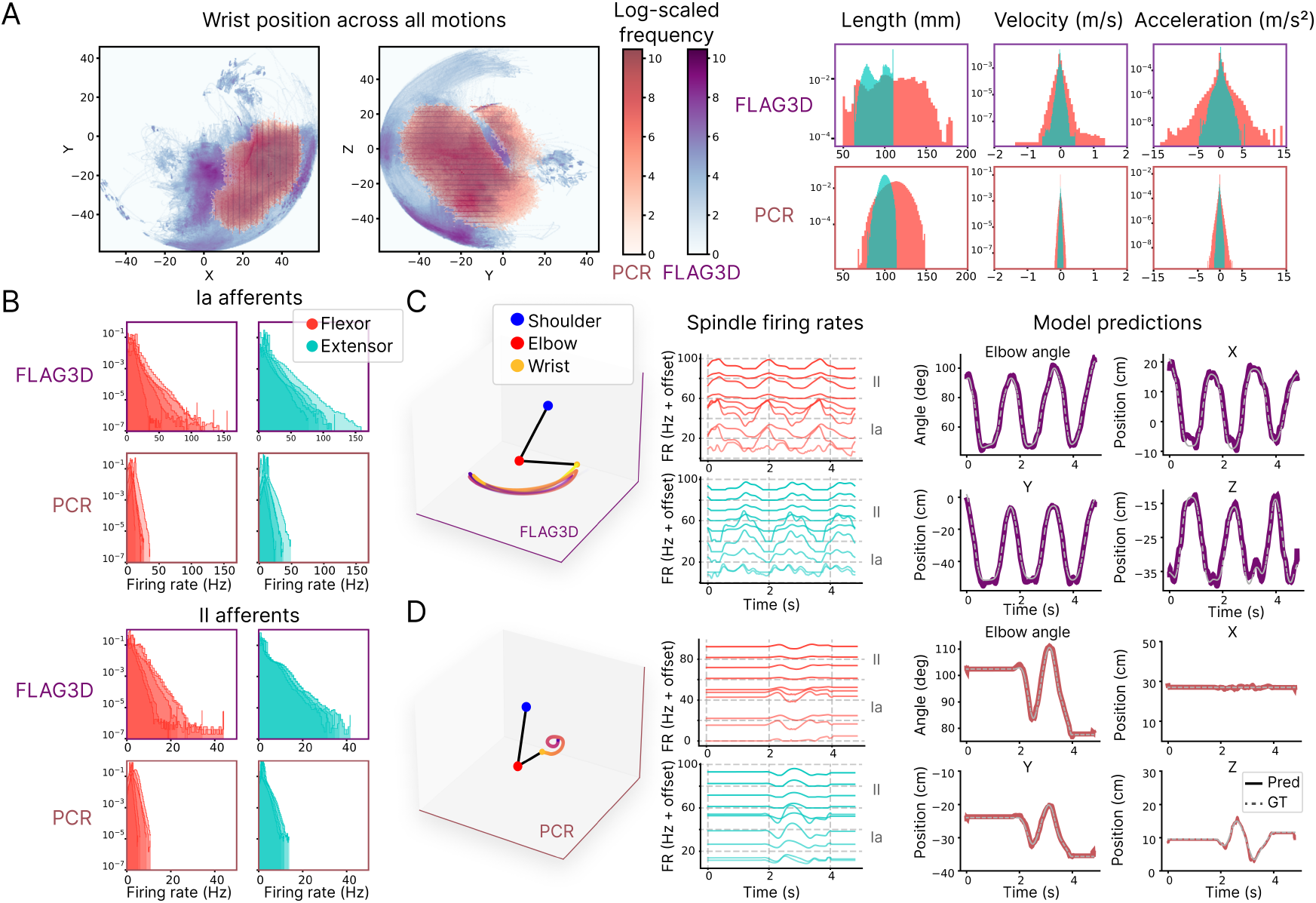
**A**. Movement statistics for FLAG3D and PCR expressed as space coverage of the wrist (left, projected on the X-Y and Y-Z planes), and distributions of muscle states (right), shown for the elbow flexor (long biceps brachii) and extensor (medial triceps brachii). FLAG3D motions cover more space compared to PCR motions (binomial test of quantized space coverage, 250 × 250 bins, *p <* 0.0001, *N*_*FLAG*3*D*_ = 5367 × 1125, *N*_*PCR*_ = 24633 × 1125 (motions × time-steps), and span a broader distribution of muscle states. **B**. The density distribution of the generated muscle spindle firing rates for Ia afferents (top) and II afferents (bottom), across FLAG3D and PCR, shown for the elbow flexor and extensor. Different lines correspond to different muscle spindles. **C**. Arm trajectories (left) generate muscle spindle signals (middle, signals are offset for visualization), which the model receives to predict the respective arm movement (right), expressed as the elbow angle (top right), the wrist position in cartesian XYZ coordinates (top right, bottom left, bottom right), and three shoulder joint angles (not shown here). Colored lines indicate the model prediction (Pred); dashed lines correspond to the ground truth (GT). Results were generated using a model trained on muscle spindle inputs from both PCR and FLAG3D (Table 3, last row). Here we only visualize spindle rates only for one flexor and extensor, but our model considers 25 arm muscles. C shows a FLAG3D test set example. **D**. Same as C, but for a PCR example.

### Modeling spindles

Of all proprioceptive sensors, we focused on muscle spindles, as those are considered the most relevant for this illusion (Marasco and de Nooij, 2023; Proske and Gandevia, 2012). Various models of muscle spindles have been developed from transfer functions and linear models (Chen and Poppele, 1978; Dimitriou and Edin, 2008; Houk et al., 1981; Poppele and Bowman, 1970; Prochazka and Gorassini, 1998) to biophysically plausible models (Blum et al., 2020; Hasan, 1983; Housley et al., 2024; Lin and Crago, 2002; Mileusnic et al., 2006; Schaafsma et al., 1991; Simha and Ting, 2024). The latter would require heavy computation to model each muscle spindle, thus we considered a transfer function approximation of the muscle spindles, which map muscle state (muscle length, velocity, and acceleration) to firing rate.

**Table 3.** Performance of various models on the test set of the PCR, FLAG3D, and elbow flexion (Ef3D) data. The first row reports the results by Sandbrink et al. (2023), the rest are PyTorch models trained with different inputs, outputs, sampling frequency, and training data. Inputs are either muscle length (l) and velocity (v) or type Ia and II afferents (N=5 per muscle and type). We report the decoding error on the wrist (x,y,z) coordinates and the RMSE on the elbow angle for different test datasets. The third row from the bottom shows the results for a single model (used for Figure 3) trained with a set of spindle parameter coefficients. The second to last row shows the mean values ± SEM for five different spindle parameter instances (N=5 models) and the last row shows the mean values ± SEM for five different spindle parameter instances and four different training instances (N=20 models)

| Model Description |  |  |  | PCR accuracy |  | FLAG3D accuracy |  | Ef3D accuracy |  |
| --- | --- | --- | --- | --- | --- | --- | --- | --- | --- |
| Freq | Train data | Inputs | Outputs | Wrist | Elbow Ang. | Wrist | Elbow Ang. | Wrist | Elbow Ang. |
| 66.7 Hz | PCR (reported) | l, v | wrist | 0.13 cm ± 0.005 | N/A | N/A | N/A | N/A | N/A |
| 66.7 Hz | PCR | l, v | wrist | 0.15 cm | N/A | N/A | N/A | 1.53 cm | N/A |
| 240 Hz | PCR | l, v | wrist, angles | 0.21 cm | 0.12° | 12.39 cm | 21.27° | 4.53 cm | 3.55° |
| 240 Hz | PCR, FLAG3D | l, v | wrist, angles | 0.22 cm | 0.16° | 0.56 cm | 0.46° | 1.35 cm | 0.56° |
| 240 Hz | PCR, FLAG3D | 5 Ia, 5 II | wrist, angles | 0.61 cm | 0.47° | 1.67 cm | 1.32° | 3.23 cm | 2.93° |
| 240 Hz | PCR, FLAG3D | 5 Ia, 5 II | wrist, angles | 0.53 cm ± 0.01 | 0.52° ± 0.02 | 1.58 cm ± 0.03 | 1.38° ± 0.04 | 3.07 cm ± 0.11 | 2.87° ± 0.16 |
| 240 Hz | PCR, FLAG3D | 5 Ia, 5 II | wrist, angles | 0.57 cm ± 0.01 | 0.56° ± 0.01 | 1.54 cm ± 0.02 | 1.42° ± 0.03 | 3.06 cm ± 0.04 | 2.84° ± 0.09 |

Following Dimitriou and Edin (2008); Prochazka and Gorassini (1998), we assumed a spindle transfer function that is either linear or includes a power term in the velocity component. In humans, Dimitriou and Edin (2008) evaluated the validity of similar transfer functions in single afferents. They found high variability in the specific coefficients (but for a limited range of movements). Instead of setting specific values for the coefficients, we arrived at a set of coefficients satisfying biological constraints on the range of the spindle firing rate for a wide range of movement statistics. Specifically, we imposed the following biological constraints on the firing rate statistics for each afferent *i* (see Methods):

- Across all arm trajectories in the movement dataset, the firing rate *f* must fall within a range [0, *f*_*max*_] Hz
- Over all the movements in the dataset, each afferent must have a certain level of lifetime activity sparseness *λ*_*s*_, defined as the proportion of time points during which the neuron has zero firing rate.

We illustrate the generated spindle data for one example of sampled parameters (Figure 2B). The generated muscle spindle data predominantly occupied sub-maximal levels, but also demonstrated a high dynamic range, which, as expected, was even more pronounced for the FLAG3D movements.

### A model of the proprioceptive pathway

We adapted the model from Sandbrink et al. (2023) in a number of ways and also re-implemented it in PyTorch (see Methods). Qualitatively, the trained models could accurately predict the wrist position and elbow angle from muscle spindle inputs both for a FLAG3D and a PCR test sample (Figure 2C,D).

We made a number of quantitative comparisons. First, for comparable settings, our PyTorch model reproduced the accuracy of the TensorFlow model from muscle length and velocity inputs (Table 3). Second, we trained the models using data sampled at a higher frequency (240 Hz, compared to the original 66.7 Hz), enabling the analysis of tendon vibration effects across a broader range of frequencies. At the higher sampling rate, the same model had similar accuracy, despite having an effectively smaller receptive field (Table 3). However, models solely trained on PCR had an average elbow accuracy of 3.55 degrees on the elbow flexion dataset (Ef3D). Third, by incorporating the FLAG3D training set, we substantially improved model accuracy across a wide physiological range, as shown in the prediction of elbow flexion movements (Ef3D). Furthermore, we extended the model’s predicted outputs to include shoulder and elbow joint angles, reflecting experimental setups where elbow angle is commonly used to measure proprioceptive illusions Roll and Vedel (1982). Finally, we refined the model inputs to represent spindle afferent firing rates rather than raw muscle state variables. This modification aligned the model more closely with the biological inputs to the proprioceptive pathway, ensuring greater physiological plausibility.

The model trained with muscle state inputs demonstrated the highest performance across all three datasets (Table 3). This outcome highlights the challenge faced by models (and the nervous system) trained with spindle inputs, as they must disentangle diverse muscle state information embedded within mixed firing rates. Despite this added complexity, spindle-based models with just five afferents per type (Ia, II) still achieved competitive performance in all three test datasets, excelling in both wrist position and elbow angle predictions (e.g., 2.9^*°*^ vs. 0.56^*°*^).

With these models in place, we asked if task-driven models are susceptible to proprioceptive illusions.

### Simulation of tendon vibration illusions

Classic experiments found that vibrations applied to the biceps tendon in an arm at rest lead to the illusion of arm extension, and vibrations applied to the triceps tendon to the illusion of arm flexion (Roll and Vedel, 1982). We simulated the tendon vibrations by perturbing the spindle firing rate based on constraints inferred from recordings of primary and secondary afferents (see Methods). While one can vary the vibration amplitude experimentally, in our case, the amplitude is implicitly set by the transfer function from vibrations to the spindle response (see Methods). We found that biceps and triceps vibrations do indeed lead to the illusion of extension and flexion. The musculoskeletal model we used included three triceps muscles representing the three heads of the muscle: the long head (tri. long), the lateral head (tri. lat.), and a medial head (tri. med.). It also included the two heads of the biceps muscle: the long (bic. long) and short (bic. short). We found that, depending on how many muscles were vibrated, the strength of the illusion varied (Figure 3A).

**Figure 3.**
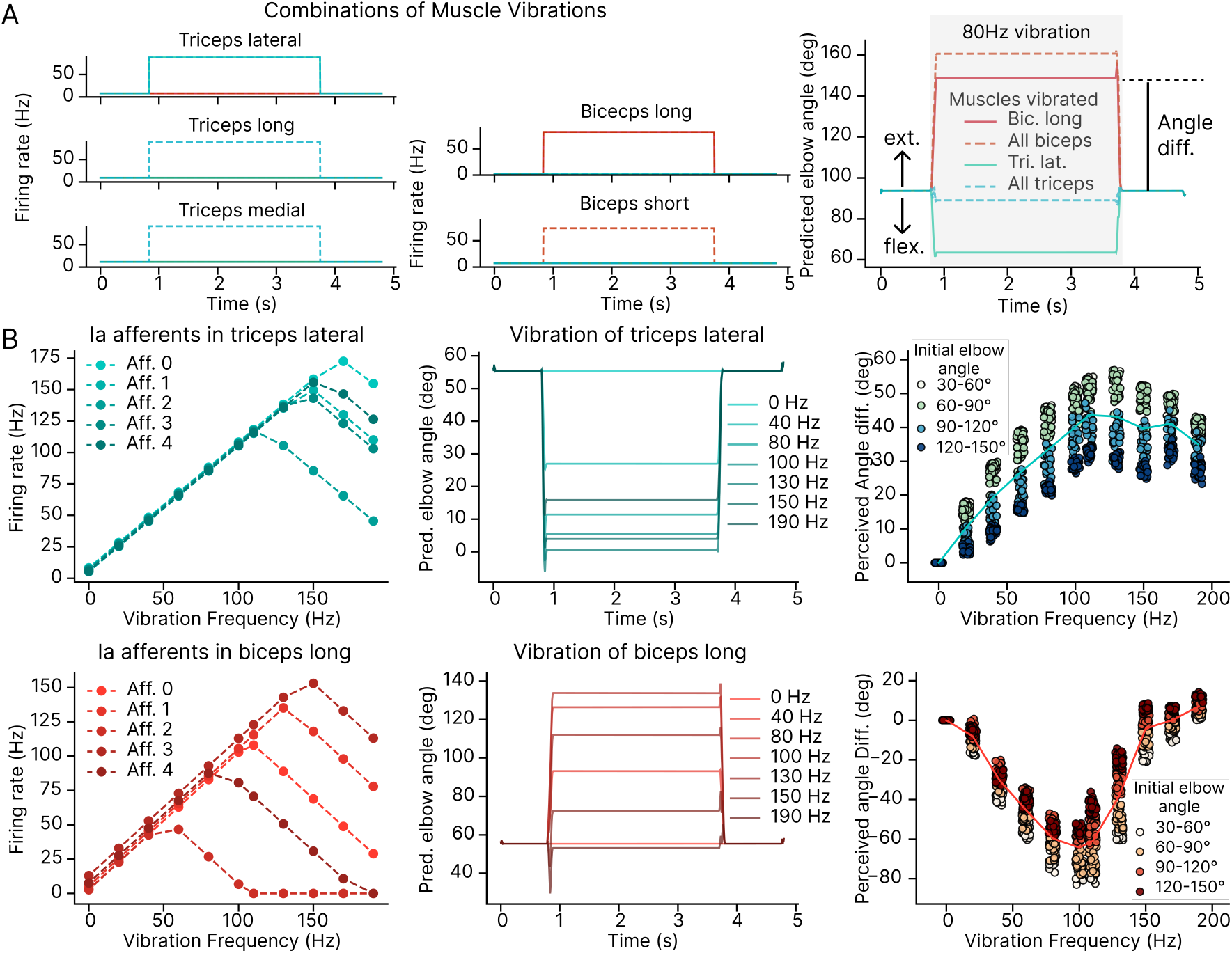
Effect of muscle-tendon vibrations on elbow angle prediction. **A**. Example demonstrating the impact of vibrating specific triceps or biceps muscle-tendons on the predicted elbow angle. The time course of the spindle discharge for one afferent is displayed, which is disrupted by the 80 Hz vibration (left). Vibrating the biceps muscles induces an illusion of extension, while vibrating the triceps muscles induces an illusion of flexion (right). **B**. Vibration frequency influences the magnitude of the proprioceptive illusion. The effect of vibration frequency on spindle discharge is modeled based on findings by Roll et al. (1989) (left). An example trial illustrates the predicted elbow angle across different vibration frequencies (middle). The measured “illusion” angle, the difference in elbow angle before and after vibration, plotted against vibration frequencies for different initial elbow angles (right, *N* = 100 per vibration frequency). Results for this figure were obtained for the model with test performance shown in the third row from the bottom in Table 3.

We systematically varied the vibration frequency (Figure 3B). At the physiological level, our synthetic Ia afferents produce a coupled response to the vibration frequency up to the maximal vibration frequency, which varies from afferent to afferent. Beyond this maximal frequency, the firing rate decreases with the vibration frequency (Figure 3B left). Thus, our modeling of the effect of vibrations on the spindle response captures the variety and richness in response found in experiments (Roll et al., 1989). At the behavioral level, our trained model reported perturbed elbow angles (Figure 3B middle, p-values in Table 4). Here, we tested 100 different arm configurations and elbow angles and found that the illusion results were consistent across different arm postures (on Es3D). The strength of the illusion depended primarily on the vibration frequency, and to a lesser extent on the initial elbow angle (Figure 3B right, p-values in Tables 5-6). Across elbow angles, we found an increase of the illusion to some maximum frequency. Beyond this point, the illusion decreased (p-values in Tables 7 and 8). This result qualitatively agrees with the data by Roll and Vedel (1982), who analyzed one particular posture.

**Table 4.**
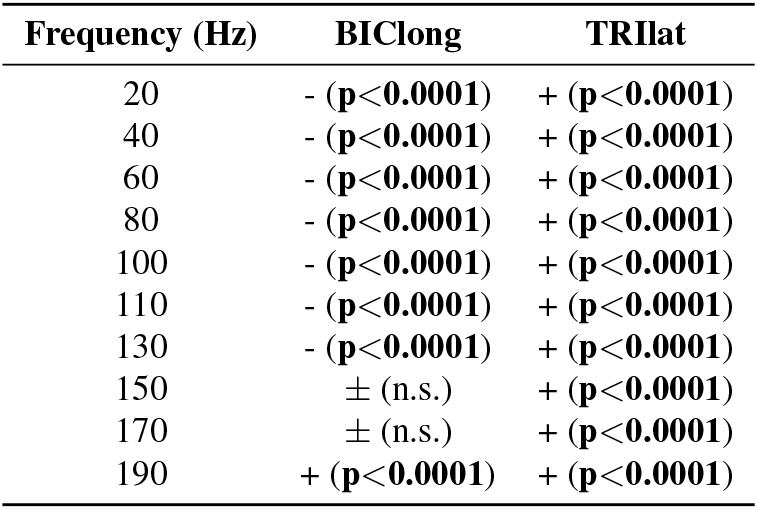
Statistical significance of the measured “illusion” angle by muscle and frequency (Figure 3B). One-sided non-parametric Wilcoxon test, *N* = 100 trials throughout. Minus (*−*) symbols indicate perceived flexion; plus (+) symbols indicate perceived extension.

**Table 5.**
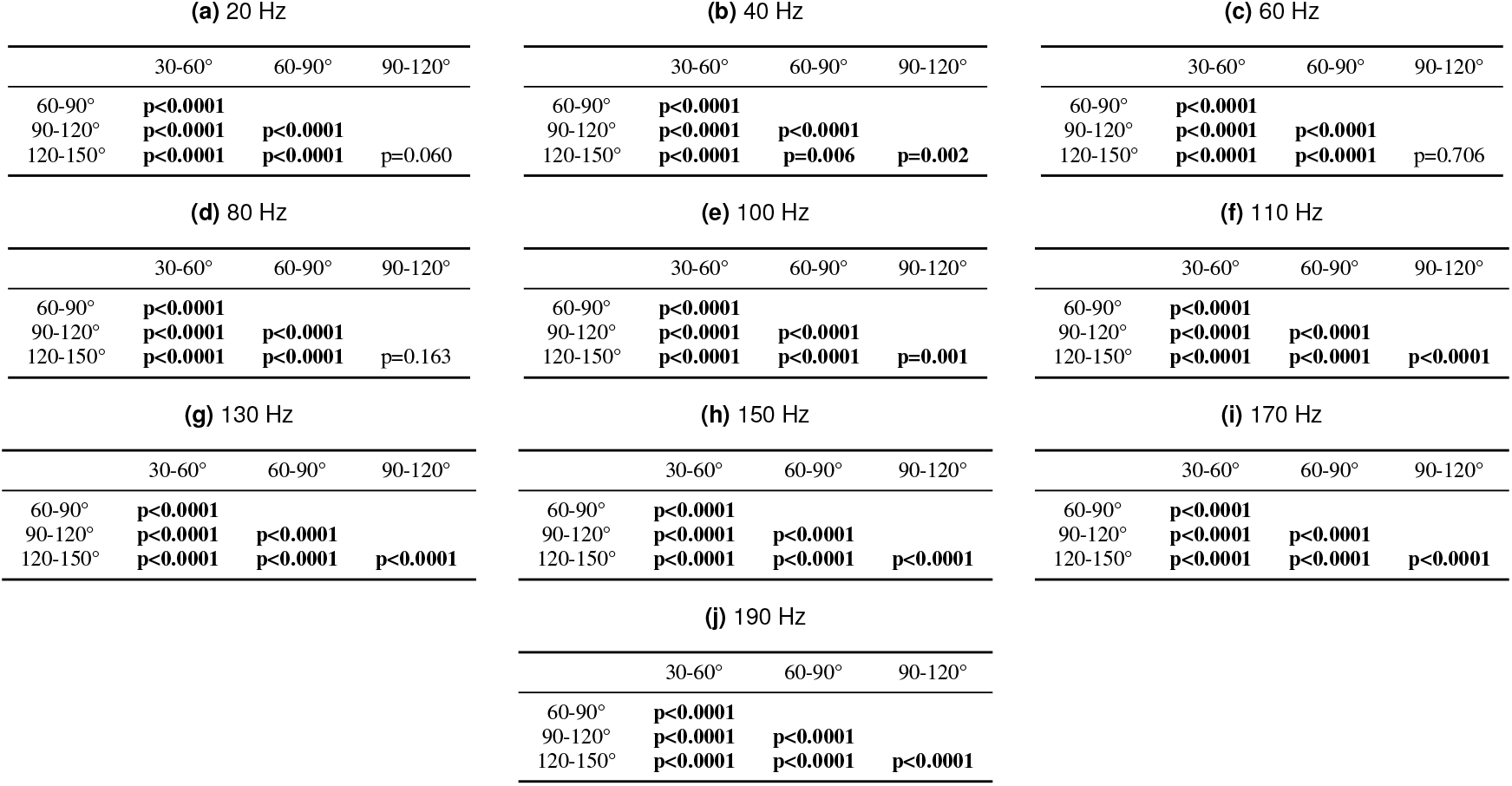
Biceps (long): Statistical differences of the measured “illusion” angle between elbow angle ranges (Figure 3B). Two-sided non-parametric Wilcoxon test. 30-60°: *N* = 17, 60-90°: *N* = 32, 90-120°: *N* = 26, 120-150°: *N* = 25.

**Table 6.**
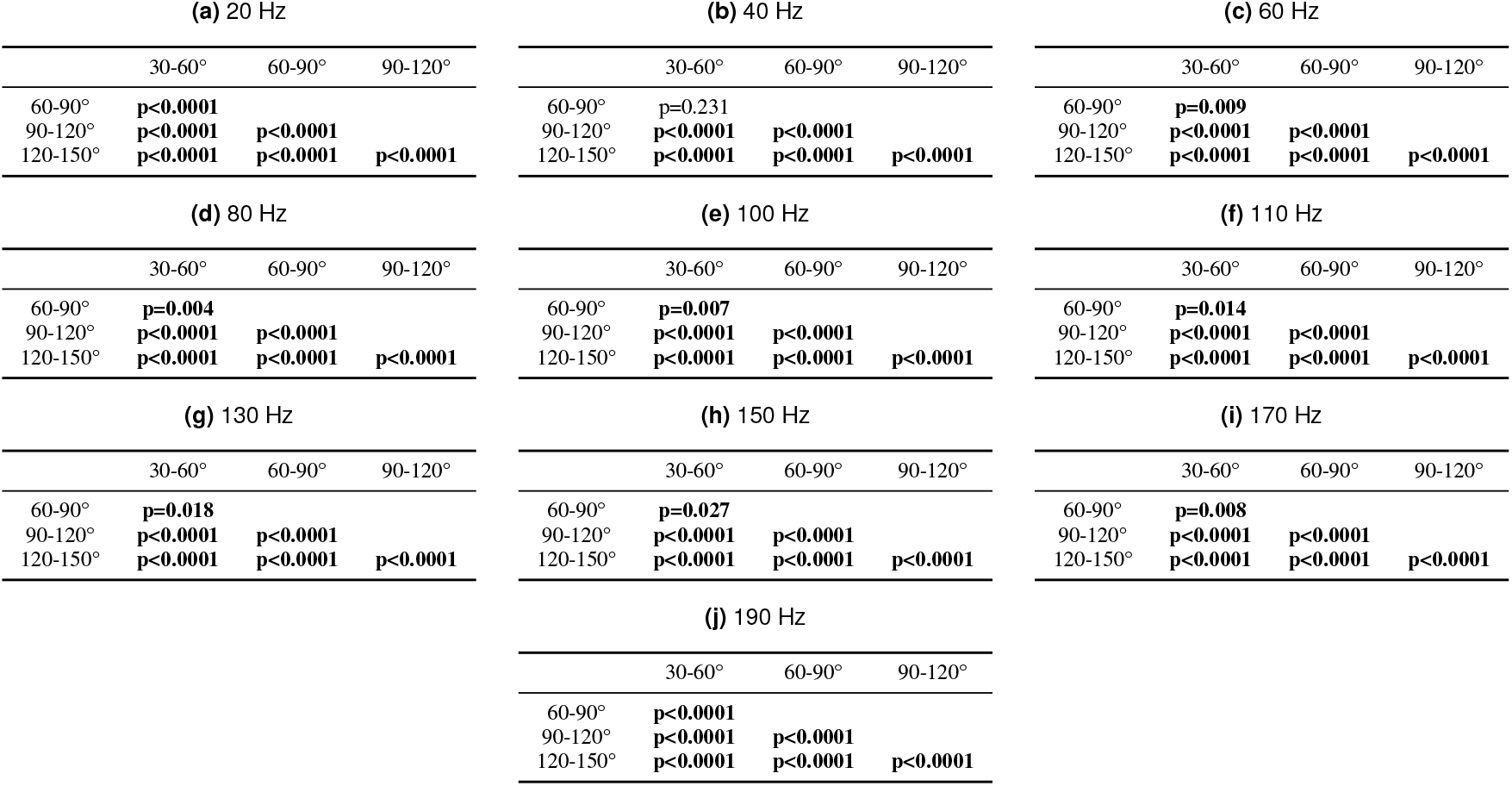
Triceps (lateral): Statistical differences of the measured “illusion” angle between elbow angle ranges (Figure 3B). Two-sided non-parametric Wilcoxon test. 30-60°: *N* = 17, 60-90°: *N* = 32, 90-120°: *N* = 26, 120-150°: *N* = 25.

**Table 7.** Biceps (long): Statistical differences of the measured “illusion” angle between vibration frequencies (Figure 3B). One-sided paired non-parametric test (Wilcoxon). *N* = 100 trials throughout.

| F2 | Frequency 1 (Hz) |  |  |  |  |  |  |  |  |
| --- | --- | --- | --- | --- | --- | --- | --- | --- | --- |
| (Hz) | 20 | 40 | 60 | 80 | 100 | 110 | 130 | 150 | 170 |
| 40 | <<br>**p<0.0001** | - | - | - | - | - | - | - | - |
| 60 | <<br>**p<0.0001** | <<br>**p<0.0001** | - | - | - | - | - | - | - |
| 80 | <<br>**p<0.0001** | <<br>**p<0.0001** | <<br>**p<0.0001** | - | - | - | - | - | - |
| 100 | <<br>**p<0.0001** | <<br>**p<0.0001** | <<br>**p<0.0001** | <<br>**p<0.0001** | - | - | - | - | - |
| 110 | <<br>**p<0.0001** | <<br>**p<0.0001** | <<br>**p<0.0001** | ±<br>n.s. | ><br>**p<0.0001** | - | - | - | - |
| 130 | <<br>**p<0.0001** | <<br>**p<0.0001** | ><br>**p<0.0001** | ><br>**p<0.0001** | ><br>**p<0.0001** | ><br>**p<0.0001** | - | - | - |
| 150 | ><br>**p<0.0001** | ><br>**p<0.0001** | ><br>**p<0.0001** | ><br>**p<0.0001** | ><br>**p<0.0001** | ><br>**p<0.0001** | ><br>**p<0.0001** | - | - |
| 170 | ><br>**p<0.0001** | ><br>**p<0.0001** | ><br>**p<0.0001** | ><br>**p<0.0001** | ><br>**p<0.0001** | ><br>**p<0.0001** | ><br>**p<0.0001** | ><br>**p<0.0001** | - |
| 190 | ><br>**p<0.0001** | ><br>**p<0.0001** | ><br>**p<0.0001** | ><br>**p<0.0001** | ><br>**p<0.0001** | ><br>**p<0.0001** | ><br>**p<0.0001** | ><br>**p<0.0001** | ><br>**p<0.0001** |

**Table 8.**
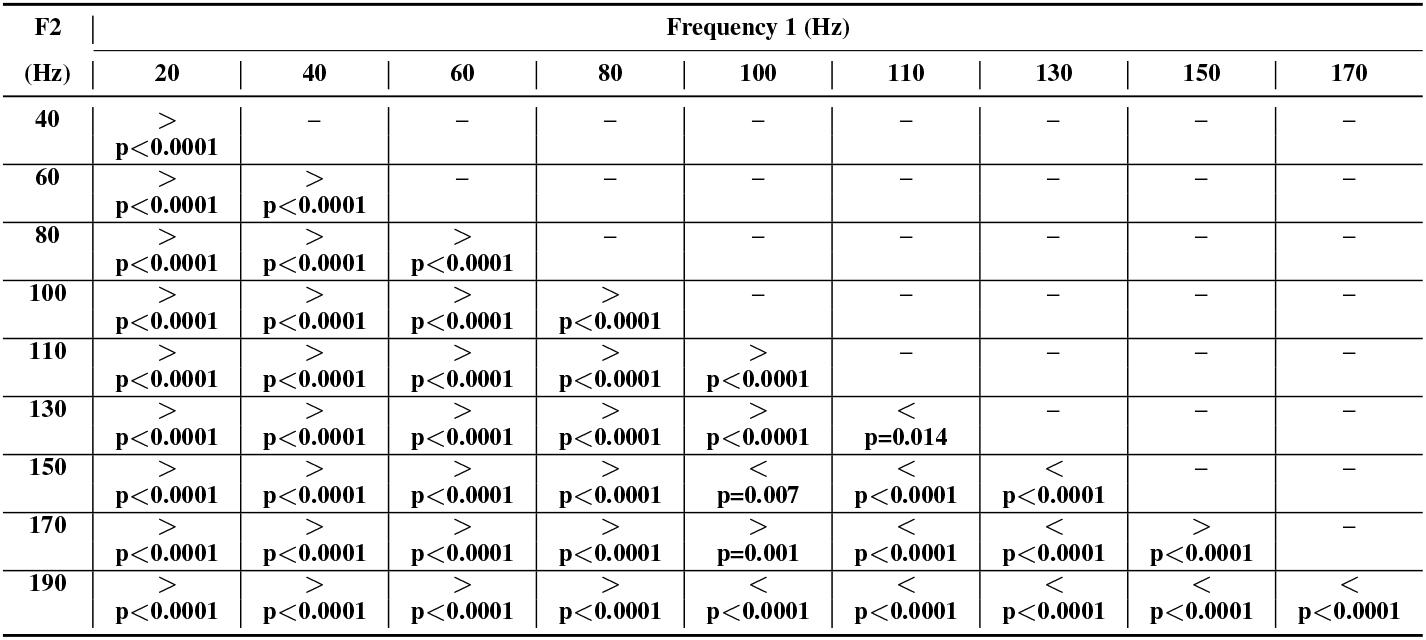
Triceps (lateral): Statistical differences of the measured “illusion” angle between vibration frequencies (Figure 3B). One-sided paired non-parametric test (Wilcoxon). *N* = 100 trials throughout.

Our results so far are based on a single model trained on one set of muscle spindle parameters. Are those results robust with respect to model and spindle parameters?

Firstly, even for the same muscle spindle parameters (input/output pairs), due to the nonconvex nature of the loss landscape for deep neural networks (Bengio et al., 2017), stochastic gradient descent might find different solutions. In other words, the learned neural network weights of two models 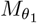 and 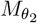 when minimizing Eq. (5) might be substantially different. Secondly, the learned models might be substantially different for different optimized muscle spindle parameters (based on Eq. (3)). From now on, we always vibrate all heads of the biceps or triceps simultaneously.

### Robustness of illusions across levels of variability

To evaluate the robustness of proprioceptive illusions to variations in spindle population responses, we sampled multiple coefficient combinations (see Methods). Five distinct coefficient sets were generated for each afferent, and four independent model instances were trained for each set. The performance of these 20 models across testing datasets demonstrated consistent accuracy regardless of spindle coefficient variations or training instances. For the PCR dataset, the models achieved an average error of 0.57 cm ± 0.01 for wrist position and 0.56° ± 0.01 for elbow angle. Similarly, on FLAG3D, the average errors were 1.54 cm ± 0.02 and 1.42° ± 0.03, while for Ef3D, the errors were 3.06 cm ± 0.04 and 2.84° ± 0.09 (last row in Table 3). We then evaluated the effect of vibrating the triceps or the biceps across different frequencies for the same 100 arm configurations across the 20 models.

Initially, we analyzed whether specific spindle coefficient configurations influenced the perceived illusion. Results showed variability in illusion strength and the frequency eliciting the maximal effect (Figure 4A). While part of the variability arose from differences across training instances, certain coefficient configurations notably altered the illusion. For some configurations, the illusion was nearly absent, while others caused a directional shift in triceps-induced illusions. Despite variability across models, encouragingly, any given model gave consistent responses for the same vibration frequency across different postures.

**Figure 4.**
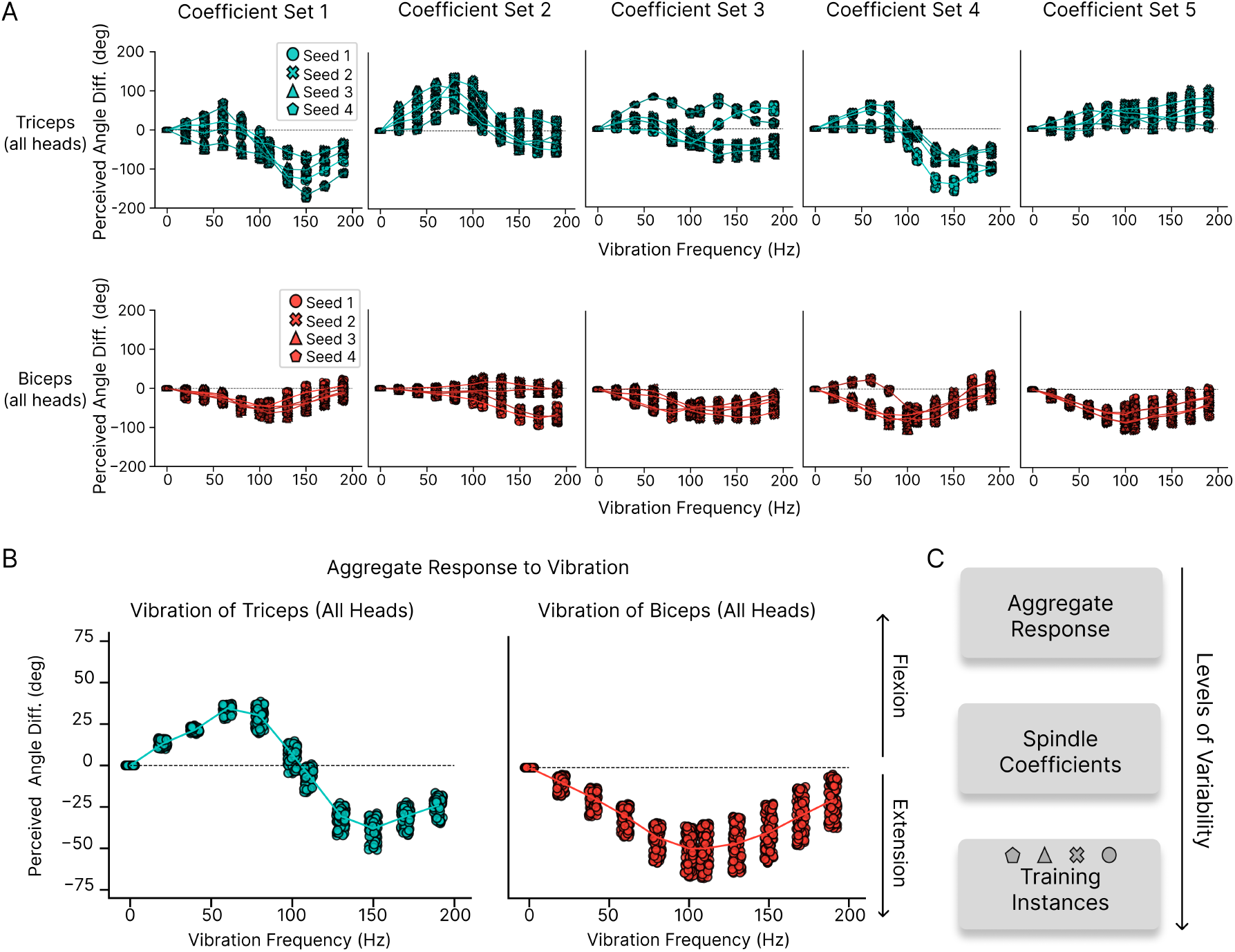
Robustness of vibration illusion **A**. Perceived angle difference for the five different spindle coefficients sets tested for vibration of the triceps (top) and biceps (bottom). Each line corresponds to the average across arm configurations for different training seeds. *N* = 100 trials per training seed, per frequency. **B**. Perceived angle difference averaged across spindle coefficient combinations and training instances for *N* = 100 different arm configurations for different vibration frequencies of all three heads of the triceps (left) and both biceps heads (right). Each dot represents the perceived angle difference for that arm configuration averaged over 20 models (5 coefficient seeds with 4 training seeds each). **C**. The variability in the illusion strength and direction increases over the choice of spindle coefficients and training instances.

Due to computational limitations when training the network models on a single GPU, each model considered just five afferents per type. This number is of course substantially smaller than the number of spindles per arm muscle (Banks, 2006; Kissane et al., 2023). Thus, “ensembling” multiple neural network models also captures the actual system better. Furthermore, when providing the vibration inputs to the models, we are driving the models with stimuli that are far away from the training distribution. In such cases, ensembling methods are widely used to find more reliable deep learning models (Ganaie et al., 2022). Thus, we averaged over seeds and spindle populations.

Averaging the predictions over all spindle coefficient combinations, and training instances, we observed a consistent effect across arm configurations (Figure 4B, p-values in Table 9). Vibrating the biceps induced the experimentally expected illusion of elbow extension, with maximal effects occurring between 100 and 110 Hz. In contrast, vibrating the triceps revealed a bimodal frequency dependence: at frequencies up to 100 Hz, the expected elbow flexion illusion occurred, but at higher frequencies, the illusion reversed, producing a prediction of elbow extension.

**Table 9.** Statistical significance of the measured “illusion” angle by muscle and frequency (Figure 4B). One-sided non-parametric Wilcoxon test, *N* = 100 trials throughout. Minus (*−*) symbols indicate perceived flexion; plus (+) symbols indicate perceived extension.

| **Vibration** | **Muscle** |  |
| --- | --- | --- |
| **Frequency (Hz)** | **Biceps (all heads)** | **Triceps (all heads)** |
| 20 | -<br>($p < 0.0001$) | +<br>($p < 0.0001$) |
| 40 | -<br>($p < 0.0001$) | +<br>($p < 0.0001$) |
| 60 | -<br>($p < 0.0001$) | +<br>($p < 0.0001$) |
| 80 | -<br>($p < 0.0001$) | +<br>($p < 0.0001$) |
| 100 | -<br>($p < 0.0001$) | +<br>($p < 0.0001$) |
| 110 | -<br>($p < 0.0001$) | -<br>($p < 0.0001$) |
| 130 | -<br>($p < 0.0001$) | -<br>($p < 0.0001$) |
| 150 | -<br>($p < 0.0001$) | -<br>($p < 0.0001$) |
| 170 | -<br>($p < 0.0001$) | -<br>($p < 0.0001$) |

### Mechanisms underlying frequency-response profiles

We next investigated the mechanisms underlying the distinct frequency-response profiles observed. Specifically, we sought to determine what factors influence the maximal perceived illusion and the vibration frequency at which this maximum occurs, and why vibrating the triceps tendon produces an illusion with maximal strength at 50 Hz while biceps vibration reaches maximum strength at 100 Hz.

To systematically address these questions, we decomposed the overall illusion into contributions from individual afferents. By applying vibration selectively to single afferents, we discovered substantial variation in illusion properties within the same muscle (Figure 5A). These variations manifested in three key dimensions: illusion strength, illusion direction, and the frequency at which maximum effect occurred. This heterogeneity primarily stemmed from the distinctive response characteristics of each afferent to vibratory stimuli.

**Figure 5.**
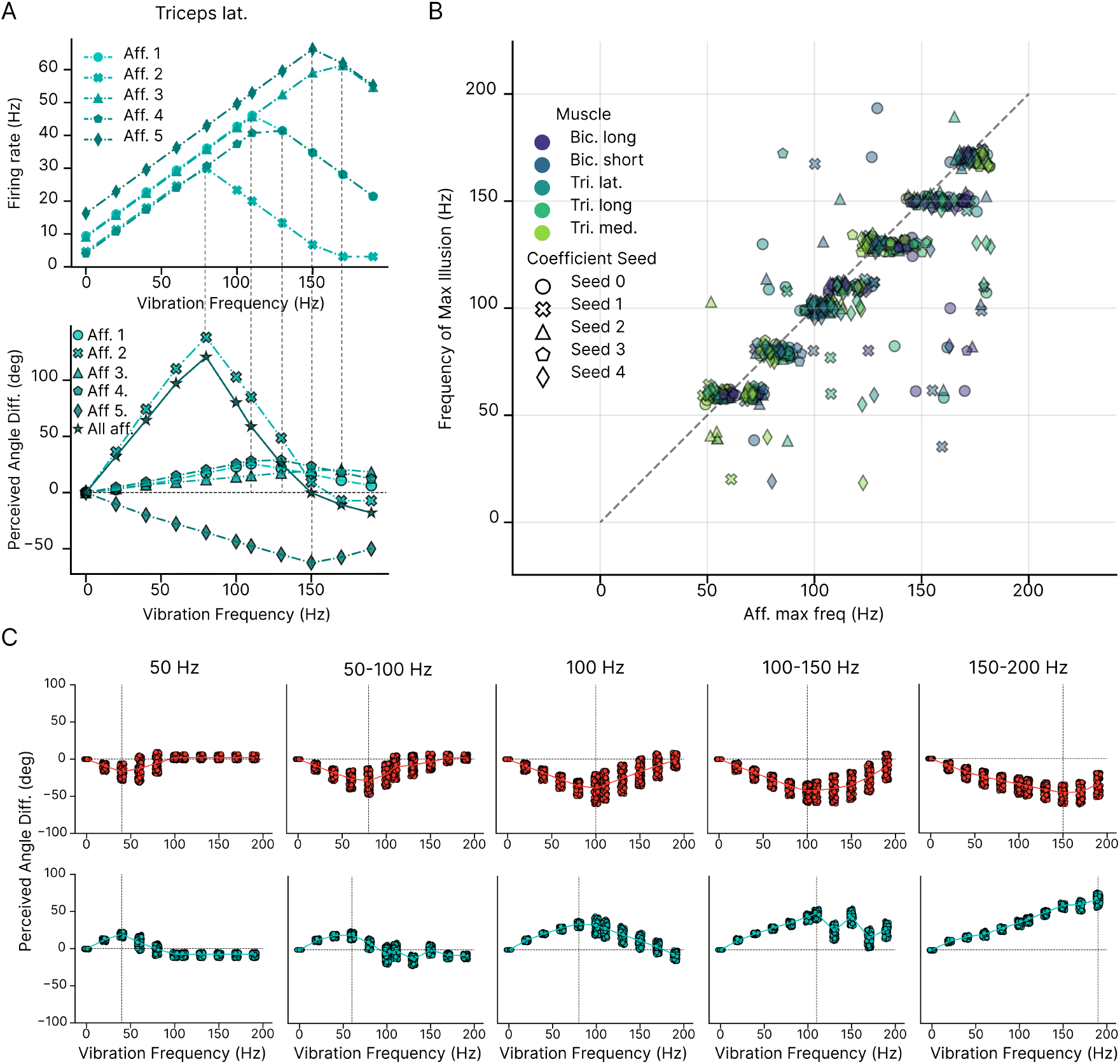
Frequency-dependent illusions **A** Individual afferents within the same muscle (triceps lateral) exhibit distinct frequency-dependent illusion profiles (bottom). This heterogeneity directly reflects differences in afferent-specific firing rate responses to vibration stimuli (top). Example of coefficient seed 1, averaged over 4 training seeds and 100 trials. **B** The frequency at which the maximum illusion occurs (*f*_*ill*_) correlates strongly with the maximum harmonic frequency (*ν*_*max*_), the maximum firing rate, of each afferent (Pearson correlation *r* = 0.86 with *p <* 0.0001). We plot 5 afferents per muscle for 5 coefficient seeds with 3 training seeds each for a total *N* = 500. **C** Perceived angle difference of vibration of all type Ia afferents in all biceps (top) and triceps muscles (bottom) as we vary the maximum harmonic frequency. We set them to a specific value (50 Hz and 100 Hz) or draw them from particular ranges (when a range is displayed). The vertical line corresponds to the frequency of maximal illusion which increases as the maximum harmonic frequency increases. Each plot represents 100 trials averaged over 20 models.

We found a strong relationship between an afferent’s maximum harmonic frequency (*ν*_*max*_), which we initially set to its maximum firing rate capacity, and the vibration frequency that elicited its peak illusion effect (*f*_*ill*_) (Figure 5B, Pearson correlation *r* = 0.86 with *p <* 0.0001). Afferents with higher maximum firing rates typically reached their peak illusion effect at higher vibration frequencies, while those with lower maximum firing rates exhibited peak effects at lower frequencies. This relationship establishes a mechanistic link between afferent physiological properties and illusion characteristics.

The maximum harmonic frequency corresponds to the highest vibration frequency at which an afferent can couple to and fire at that frequency. Roll et al. (1989) found that this maximum frequency could vary between 40 and 180 Hz across type Ia afferents from ankle muscles (tibialis anterior and extensor digitorium longus), but the distribution in different arm muscles remains unclear. To explore this, we investigated how modifying the maximum harmonic frequency affects the frequency dependence of illusions when vibrating all type Ia afferents in either triceps or biceps muscles.

We had initially taken the maximum harmonic frequency to correspond to the maximum firing rate for each afferent. Setting the maximum harmonic frequency *ν*_*max*_ to 100 Hz for all afferents affected illusions in the biceps muscles minimally, but significantly altered the triceps response, rendering it unimodal (Figure 5C). This adjustment shifted the maximum illusion frequency to 100 Hz and generated an illusion of flexion across all frequencies up to 190 Hz.

When we allowed the maximum harmonic frequency to vary across afferents within increasing ranges (50 Hz, 50-100 Hz, 100-150 Hz, 150-200 Hz), we found that for both triceps and biceps, the frequency of maximum illusion shifted correspondingly with the range of maximum harmonic frequency (Figure 5C). Notably, when all afferents’ maximum harmonic frequencies were drawn from these tighter intervals, we did not observe the clear reversal of illusion direction seen for the triceps (Figure 4B).

This finding suggests that the reversal in illusion direction requires tendon vibration to engage multiple muscles containing afferents with a broad range of harmonic frequencies. We would expect such reversals to be less likely in muscles with biological levels of afferent density (over 100 afferents per muscle Banks (2006); Kissane et al. (2023)), as the diverse frequency responses would likely average out across the population, diminishing the probability of directional shifts in the perceived illusion.

These results highlight the relationship between peripheral physiological properties of muscle spindles (maximum harmonic frequency) and central perceptual illusions. Furthermore, they link the frequency of maximum illusion with afferent firing rate statistics, suggesting that strong illusions at higher vibration frequencies require muscle afferents capable of responding to those frequencies.

### Effect of vibrating secondary afferents on illusions

We initially modeled tendon vibration as affecting only type Ia afferents, based on experimental measurements of type Ia and type II afferent firing rates during low-amplitude tendon vibrations(Roll et al., 1989). However, type II afferents can show more sensitivity to high-amplitude vibrations (Burke et al., 1976). Here, we investigate how the vibration of type II afferents influences the perceived illusions.

Primary and secondary afferents differ in their physiological response properties: secondary afferents are more sensitive to static muscle length changes, while primary afferents predominantly detect dynamic changes (Banks et al., 2021; Matthews and Stein, 1969). Secondary afferents also typically exhibit lower firing rates. Our model captured these physiological differences through distinct spindle parameter ranges for primary and secondary afferents (Figure 6A).

**Figure 6.**
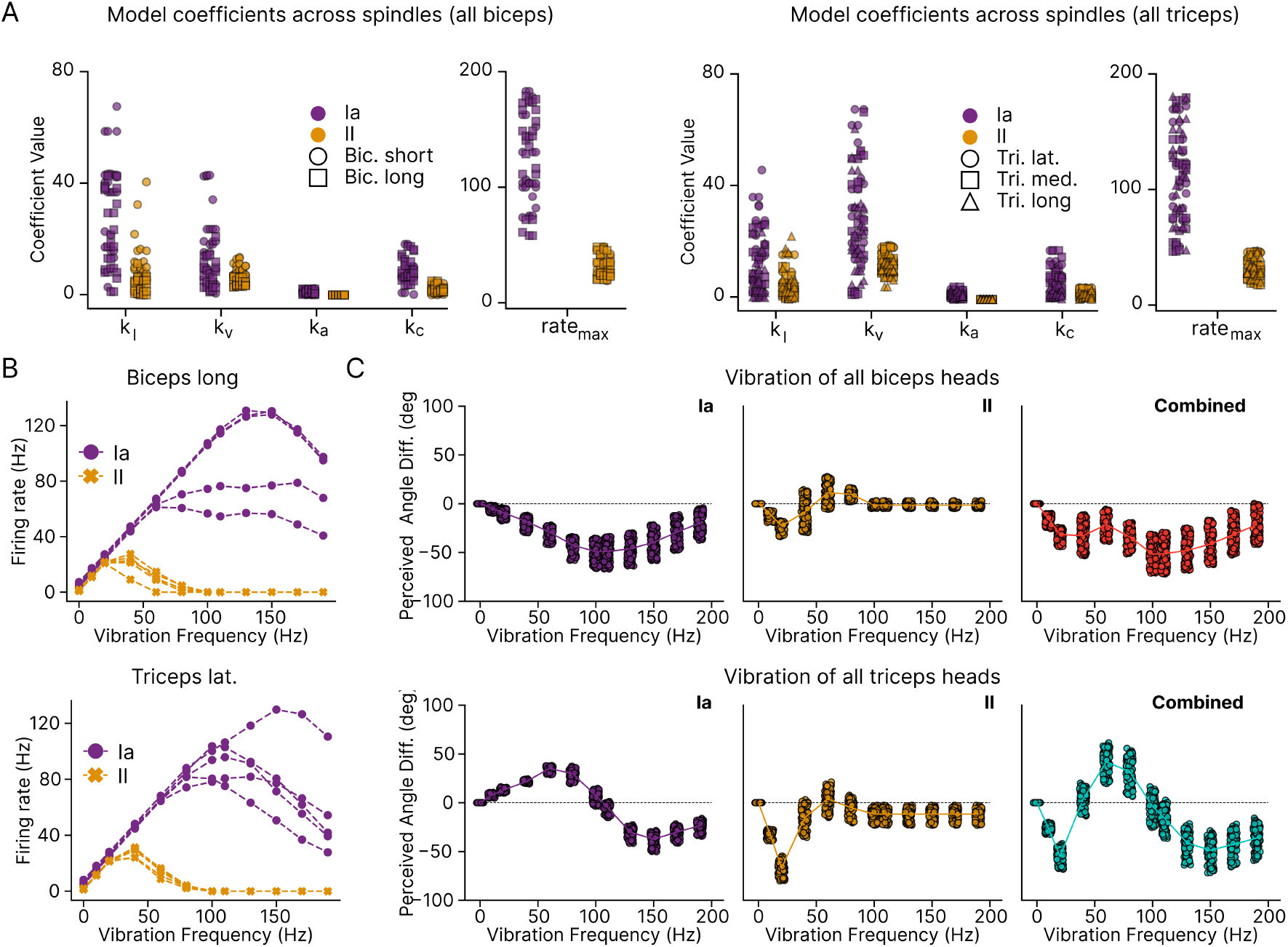
Effect from vibrating type II afferents. **A** Spindle coefficients for type Ia and type II afferents for both biceps heads (left) and all three triceps heads (right), computed by optimizing the defined cost function (see Methods). Type Ia afferents exhibit higher maximum firing rates and larger coefficients *k*_*i*_ for muscle length, velocity, acceleration, and baseline firing rate compared to type II afferents. These differences reflect the greater sensitivity of type Ia afferents to dynamic changes in muscle stretch, whereas type II afferents primarily respond to static length. (Sample size: *N* = 25 for each afferent type and each muscle corresponding to the 5 different afferent per coefficient seed). **B** Effect of vibration on afferent firing rates. Increasing vibration frequency differentially affects the firing rates of type Ia and type II afferents in the biceps long (top) and triceps lateral (bottom) (5 type Ia and 5 type II afferents represented). **C** Perceived illusions induced by vibration. Vibration applied selectively to type Ia afferents (left), type II afferents (middle), or both combined (right) alters perceived elbow angle for all biceps (top) and all triceps (bottom) muscles. (*N* = 100 trials per vibration frequency averaged over 20 models (5 coefficient seeds with 4 training seeds each) as in Figure 4B).

Previous work by Roll et al. (1989) demonstrated that few secondary afferents responded to tendon vibrations, those that did, showed a lower harmonic vibration frequency compared to primary afferents. Consistent with these findings, in our model, type II afferents reach maximal firing rate responses at lower vibration frequencies (Figure 6B).

When examining illusion generation, we found that vibrating type II afferents alone produced illusions only at low vibration frequencies (below 100 Hz) (Figure 6C). The maximal illusion for type II afferent stimulation occurred at lower vibration frequencies (20 Hz) compared to type Ia afferent stimulation. These results also corroborate our observation about the tight link between the maximal harmonic frequency of afferents and the perceptual frequency dependency.

When both afferent types were vibrated simultaneously, the resulting illusion was predominantly determined by type II afferents at low frequencies but followed the pattern of type Ia afferents at higher frequencies. These results indicate that type Ia afferents are necessary for generating illusions at higher vibration frequencies, consistent with experimental observations.

### Muscle-specific effects on illusion strength

Our computational model and simulation framework allowed us to systematically investigate the effects of vibrating each of the 25 modeled muscles (Table 2). We observed muscle-specific differences in the resulting perceived elbow angle. Vibration of some muscles produced robust biases in the perceived elbow angle, while others generated minimal perceptual effects (Figure 7). For example, the vibration of brachialis, a muscle involved in the elbow flexion, induced an extension illusion similar to that observed with biceps stimulation. Conversely, vibration of anconeus, a muscle involved in elbow extension, produced a flexion illusion resembling the effect of triceps vibration. In contrast, vibration of the thoracic latissimus dorsi, involved in shoulder movement, elicited negligible changes in perceived elbow angle (Figure 7A).

**Figure 7.**
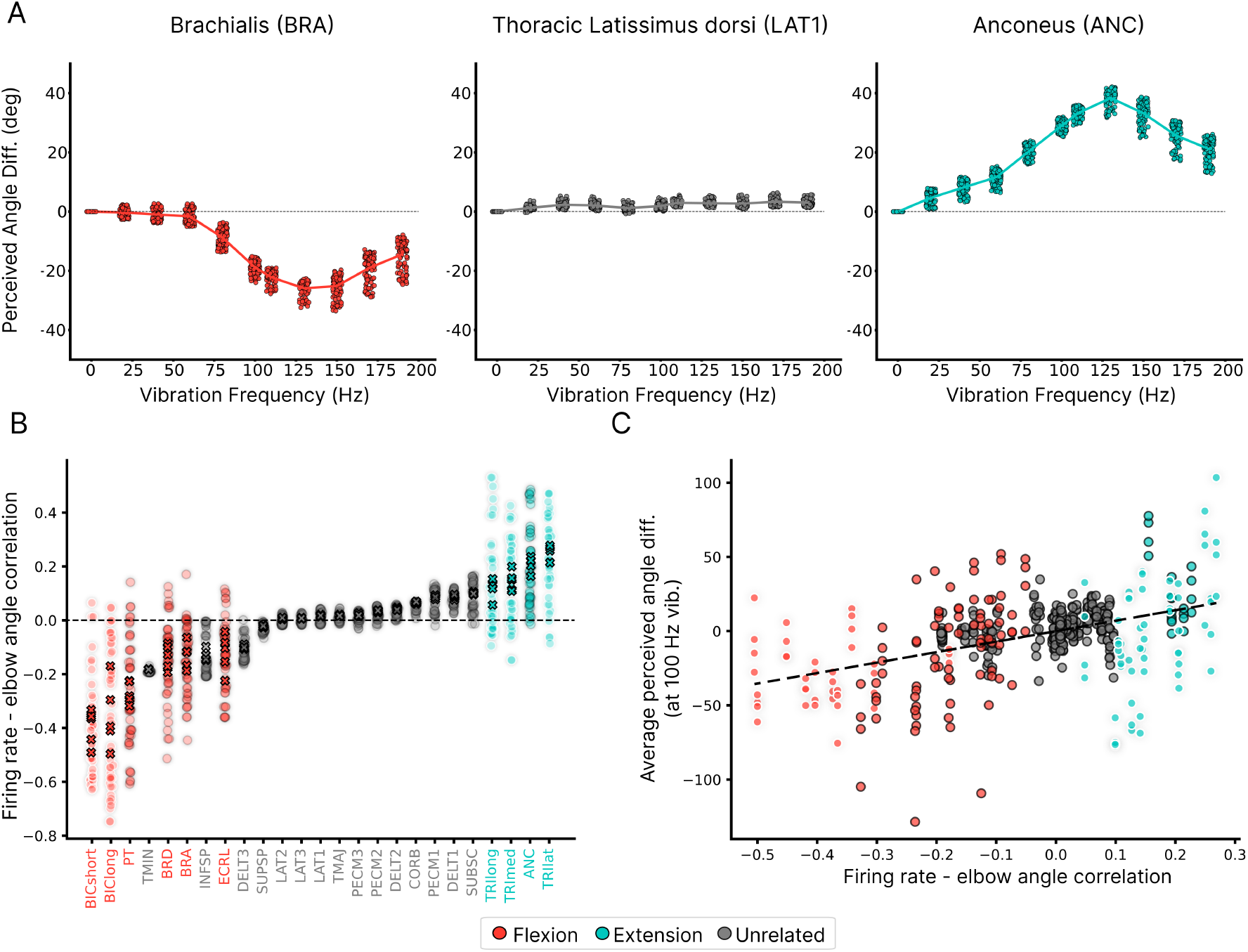
Effect from vibrating other muscles. **A**. Frequency-dependent illusion profiles for three representative muscles beyond the standard biceps and triceps. Brachialis vibration (left) produces an extension illusion similar to biceps, anconeus vibration (right) produces a flexion illusion similar to triceps, while thoracic latissimus dorsi vibration (middle) generates minimal perceptual effects. Each data point represents the mean perceived elbow angle difference across 100 trials, averaged across all trained models. **B**. Pearson correlation coefficients between afferent firing rates and elbow flexion angle across the training dataset for each modeled upper arm muscle. Flexion-related muscles exhibit strong negative correlations (increased firing with flexion), extension-related muscles show strong positive correlations, and muscles with minimal involvement in elbow movement display correlations near zero (*N* = 10 afferents per muscle for each of the five coefficient sets). Crosses indicate the mean correlation across afferents for each coefficient set. **C**. Relationship between the afferent-angle correlation (averaged across 10 afferents for each of the five coefficient sets) and mean vibration-induced illusion across 100 trials (five trained models per coefficient set) at 100 Hz. The value of the correlation coefficient correlates significantly with the strength of the proprioceptive illusions (Pearson *r*^2^ = 0.2, *p <* 0.0001.)

These differential responses indicated that our model successfully captured the functional relationships between specific muscles and elbow angle perception. To quantify this relationship, we calculated the mean correlation between afferent responses and elbow angle across the training dataset for each muscle (Figure 7B). This analysis revealed that the statistical patterns in our training data reflected the known functional roles of different muscles. Afferents from muscles associated with elbow flexion (biceps, pronator teres, brachioradialis, and brachialis) displayed strong negative correlations with elbow angle, indicating that increased firing rates in these afferents corresponded to decreasing elbow angles (greater flexion). Conversely, afferents from extension-related muscles (triceps and anconeus) exhibited strong positive correlations.

The magnitude of this correlation coefficient (regardless of direction) proved to be a powerful predictor of illusion strength during 100 Hz vibration. Muscles with stronger correlations (either positive or negative) consistently produced more pronounced illusions, while those with correlations closer to zero generated minimal perceptual effects (Figure 7C). This relationship establishes a direct link between a muscle’s functional role in joint movement and its capacity to induce proprioceptive illusions when vibrated.

## Discussion

Vibrations applied to muscle-tendons are well-known to induce various proprioceptive illusions (Eklund, 1972; Goodwin et al., 1972a,b; Roll and Vedel, 1982). Here, we developed a simple perceptual model to investigate whether feedforward models of the ascending proprioceptve pathway, trained on inputs from Ia and II fibers to localize the body’s state, are also susceptible to such illusions. Our findings reveal that tendon vibrations, which drive afferent responses when processed through the optimized proprioceptive pathway, lead to perceptual illusions, suggesting parallels between our model and human proprioceptive phenomena.

Interestingly, we found variability in the perceptual illusions across models with respect to different spindle sets, and training. For some experiments, there is also substantial intersubject variability in illusion strength (Goodwin et al., 1972a; Roll and Vedel, 1982). In some cases, participants were excluded due to a lack of experienced illusion (Roll et al., 2009). An important open question is whether the variability observed in our modeling framework reflects the variability seen in experimental studies. We believe that our models provide a valuable tool for investigating how spindle encoding influences proprioceptive illusions, even if each individual model includes only ten representative afferents per muscle, whereas human muscles can contain hundreds of spindles. While computational constraints currently limit the feasibility of modeling a larger number of afferents, it would be interesting to assess whether increasing the number of afferents impacts the robustness and strength of vibration-induced effects. However, we considered a more realistic (in terms of number of spindles and robustness) model by model averaging. Indeed, by ensembling multiple trained models, we found that vibrating the biceps induced the experimentally expected illusion of elbow extension, with maximal effects occurring between 100 and 110 Hz. In contrast, vibrating the triceps revealed a bimodal frequency dependence: at frequencies up to 100 Hz, the expected elbow flexion illusion occurred as expected, but at higher frequencies, the illusion reversed, producing a prediction of elbow extension.

Digging into the mechanism, we found a dependency between the peripheral physiological properties of muscle spindles—specifically, an afferent’s maximal sensitivity to tendon vibrations—and the frequency dependence of central perceptual illusions. This relationship suggests that variations in spindle dynamics could directly shape the perception of limb position and movement, reinforcing the idea that proprioceptive illusions are not purely a central phenomenon but are tightly linked to peripheral encoding mechanisms. These findings highlight the need for a more detailed characterization of afferent responses to vibration across different muscles and individuals. Systematic, large-scale measurements of afferent tuning properties could provide deeper insight into how peripheral factors influence proprioceptive perception.

Our computational model revealed a significant relationship between initial arm configuration and the magnitude of vibration-induced illusions. Previous models of proprioceptive encoding for illusions Bergenheim et al. (2000); Ribot-Ciscar et al. (2002) based on population decoding approaches did not exhibit this position-dependency. Future psychophysics studies should specifically quantify how illusion strength varies across different joint angles, which would provide valuable validation of our model’s predictions. Such experiments could systematically map the relationship between starting posture and perceptual outcomes. This would not only validate our computational approach but could also establish a more comprehensive framework for understanding how the brain interprets afferent signals in different postural contexts.

While our model did not explicitly parameterize vibration amplitude, we examined distinct scenarios that effectively capture different amplitude conditions. Under low-amplitude vibration conditions, we modeled selective activation of type Ia afferents, whereas high-amplitude conditions incorporated activation of both type Ia and type II afferents. This distinction revealed a frequency-dependent contribution pattern: type Ia afferents predominantly drive illusions at higher vibration frequencies (above 50 Hz), while type II afferents significantly influence the direction and magnitude of illusions at lower frequencies (below 50 Hz). This finding suggests that experimental manipulations targeting different vibration amplitudes and frequencies could selectively engage different proprioceptive processing pathways, offering a potential method for dissociating the contributions of primary and secondary afferents in human proprioception.

This work contributes to the growing body of research on goal-driven models for both behavioral and neural tasks (Kanwisher et al., 2023; Kar and DiCarlo, 2024; Mathis et al., 2024). In contrast to our earlier studies of the proprioceptive pathway that primarily investigated neural coding (Sandbrink et al., 2023; Vargas et al., 2024), our focus shifted towards perception, which presents a critical test for such models. Perceptual proprioceptive illusions, in particular, offer a compelling framework for studying the mechanisms and limits of conscious limb position perception as well as evaluating whether task optimized models can capture the nuances of proprioceptive processing. Importantly, our models were not explicitly trained to exhibit illusions, but rather to transform proprioceptive inputs to achieve hypothesized goals of proprioception (Sandbrink et al., 2023; Vargas et al., 2024). Nonetheless, they naturally demonstrated susceptibility to perceptual distortions under conditions of vibration. This result underscores the utility of task-optimized models in replicating the emergence of complex proprioceptive phenomena.

In contrast to classic Bayesian models for visual illusions (Geisler and Kersten, 2002; Gregory, 1968; Weiss et al., 2002), our approach employed a supervised learning paradigm. Specifically, we trained the ascending proprioceptive pathway to estimate the state of the body based on the statistics of muscle spindle inputs in the absence of vibrations. Inspired by the concept of cross-modal calibration (Sandbrink et al., 2023), we propose that visual or motor systems could provide the necessary supervisory signal to tune the proprioceptive pathway. By leveraging this framework, we bypass the need for explicit prior assumptions, instead relying on a data-driven framework of the sensory pathway. A key feature of our approach is the ability to directly read out the prediction of the model (or ensemble models) without relying on reporting mechanisms such as bimanual matching. As we are not modeling integration times and response latencies, our model practically instantly reports a shift in the perceived elbow angle upon application of muscle-tendon vibrations. In contrast, subjects report their illusionary motion percept relatively slowly with a delay (Chancel et al., 2018; Roll and Vedel, 1982). In future work, we will include those features and ask if the reaction times are predicted from physiological delays. Importantly, reporting the subjective experience via a motor action likely adds further delays. Thus, by analyzing neural representations along the proprioceptive pathway (Vargas et al., 2024) we could study what part of the reaction time is due to the delayed subjective experience vs. the readout mechanism.

Our study has several limitations. First, we focused on sensory models trained exclusively on passive data, examining whether this setting is sufficient to reproduce proprioceptive illusions. While our approach isolates key aspects of proprioceptive processing, it does not account for active states of muscles. Empirical evidence indicates that proprioceptive organs exhibit different responses to tendon vibrations depending on whether the muscles are resting (Burke et al., 1976) or contracting (Fallon and Macefield, 2007). Additionally, other sensors might play a role in these illusions (Collins et al., 2005; Proske and Gandevia, 2012). Furthermore, the computational model for muscle spindles was deliberately chosen to be simple due to the lack of (human) models for diverse behaviors. This raises an intriguing question: what are the fundamental characteristics of muscle spindle activity in terms of population firing rate statistics? This question could be addressed with emerging population recording techniques for muscle spindles.

Goodwin et al. surmised that biased perception may result from subcortical control circuits (Goodwin et al., 1972b,b). By focusing solely on the ascending proprioceptive pathway and omitting efference copies and subcortical circuits, our model demonstrates that higher-level features from subcortical or cortical regions are not strictly necessary to reproduce some proprioceptive illusions. Future work should incorporate hierarchical circuits to investigate the contributions of different neural loops to proprioceptive illusions, for instance by considering volitional control mechanisms and alpha-gamma collateral modulation of spinal reflexes (Banks, 2024; Niyo et al., 2024), more realistic spindle models (Dimitriou, 2022; Mileusnic et al., 2006; Perez Rotondo et al., 2024) and visual input that has been shown to influence illusory limb position and ownership (Blanke et al., 2015; Botvinick and Cohen, 1998; Palluel et al., 2011). Advances in learning controllers and biomechanical simulators offer exciting opportunities towards these goals (Caggiano et al., 2022; Chiappa et al., 2024).

Lastly, another limitation lies in the lack of large-scale behavioral validation. Here we only qualitatively compared our findings to classic behavioral studies (Roll and Vedel, 1982), but future research should leverage high-throughput methodologies combining virtual reality, robotics (Albert et al., 2024), and motion capture (Mathis et al., 2020), to provide quantitative data for model training and validation. Indeed, the nature of in silico models afforded us the ability to systematically test a broad range of arm configurations, a feat which has not yet been attempted experimentally. This presents an opportunity to initiate a virtuous cycle of hypothesis generation and empirical testing regarding the effect of limb position on kinesthetic perception. Since it is not feasible to conduct all necessary human experiments in vivo, in silico models can identify key effects, guiding the design of targeted in vivo studies to validate and refine our understanding of proprioceptive processing.

## Acknowledgments

We thank Alberto Chiappa and Andy Bonnetto for helpful technical input. We thank Michael Dimitriou, Anne Kavounoudias, Bianca Ziliotto and Alessandro Marin Vargas for discussions.

## Author contributions

A.M. conceived the project. M.S. generated the spindle data. A.P.R., M.S., F.D. and S.P. wrote the modeling code and analyzed the data. A.P.R. and M.S. ran the experiments and made the figures. All authors interpreted the data. A.P.R., and A.M. wrote the manuscript with input from all authors. A.M. supervised the project and acquired funding.

## Footnotes

1 The word channel in the machine learning sense, denoting certain input dimensions (Paszke et al., 2019).

## References

Albert, L., Potheegadoo, J., Herbelin, B., Bernasconi, F., and Blanke, O. (2024). Numerosity estimation of virtual humans as a digital-robotic marker for hallucinations in parkinson’s disease. Nature Communications, 15(1):1905. 10.1038/s41467-024-45912-w.

Banks, R. W. (2006). An allometric analysis of the number of muscle spindles in mammalian skeletal muscles. Journal of Anatomy, 208(6):753–768. 10.1111/j.1469-7580.2006.00558.x.

Banks, R. W. (2024). There and back again: 50 years of wandering through terra incognita fusorum. Experimental Physiology, 109(1):6–16. DOI: 10.1113/EP090760.

Banks, R. W., Ellaway, P. H., Prochazka, A., and Proske, U. (2021). Secondary endings of muscle spindles: Structure, reflex action, role in motor control and proprioception. Experimental Physiology, 106(12):2339–2366. DOI:10.1113/EP089826.

Bengio, Y., Goodfellow, I., and Courville, A. (2017). Deep learning, volume 1. MIT press Cambridge, MA, USA.

Bergenheim, M., Ribot-Ciscar, E., and Roll, J.-P. (2000). Proprioceptive population coding of two-dimensional limb movements in humans: I. Muscle spindle feedback during spatially oriented movements. Experimental Brain Research, 134(3):301–310. DOI: 10.1007/s002210000471.

Blanke, O., Slater, M., and Serino, A. (2015). Behavioral, neural, and computational principles of bodily self-consciousness. Neuron, 88(1):145–166. 10.1016/j.neuron.2015.09.029.

Blum, K. P., Campbell, K. S., Horslen, B. C., Nardelli, P., Housley, S. N., Cope, T. C., and Ting, L. H. (2020). Diverse and complex muscle spindle afferent firing properties emerge from multiscale muscle mechanics. eLife, 9:e55177. 10.5061/dryad.vdncjsxsw.

Botvinick, M. and Cohen, J. (1998). Rubber hands ‘feel’touch that eyes see. Nature, 391(6669):756–756. 10.1038/35784.

Burke, D., Hagbarth, K.-E., Löfstedt, L., and Wallin, B. (1976). The responses of human muscle spindle endings to vibration of non-contracting muscles. The Journal of physiology, 261(3):673–693. 10.1113/jphysiol.1976.sp011580.

Caggiano, V., Wang, H., Durandau, G., Sartori, M., and Kumar, V. (2022). MyoSuite – a contact-rich simulation suite for musculoskeletal motor control.

Chancel, M., Landelle, C., Blanchard, C., Felician, O., Guerraz, M., and Kavounoudias, A. (2018). Hand movement illusions show changes in sensory reliance and preservation of multisensory integration with age for kinaesthesia. Neuropsychologia, 119:45–58. 10.1016/j.neuropsychologia.2018.07.027.

Chen, W. and Poppele, R. E. (1978). Small-signal analysis of response of mammalian muscle spindles with fusimotor stimulation and a comparison with large-signal responses. Journal of Neurophysiology, 41(1):15–27. 10.1152/jn.1978.41.1.15.

Chiappa, A. S., Tano, P., Patel, N., Ingster, A., Pouget, A., and Mathis, A. (2024). Acquiring musculoskeletal skills with curriculum-based reinforcement learning. Neuron, 112(23):3969–3983.e5. 10.1016/j.neuron.2024.09.002.

Collins, D. F., Refshauge, K. M., Todd, G., and Gandevia, S. C. (2005). Cutaneous receptors contribute to kinesthesia at the index finger, elbow, and knee. Journal of Neurophysiology, 94(3):1699–1706. 10.1152/jn.00191.2005.

Dayan, P., Hinton, G. E., Neal, R. M., and Zemel, R. S. (1995). The helmholtz machine. Neural Computation, 7(5):889–904. 10.1162/neco.1995.7.5.889.

Delp, S. L., Anderson, F. C., Arnold, A. S., Loan, P., Habib, A., John, C. T., Guendelman, E., and Thelen, D. G. (2007). Opensim: open-source software to create and analyze dynamic simulations of movement. IEEE Transactions on Biomedical Engineering, 54(11):1940–1950. 10.1109/TBME.2007.901024.

Dimitriou, M. (2022). Human muscle spindles are wired to function as controllable signal-processing devices. eLife, 11:e78091. 10.7554/eLife.78091.

Dimitriou, M. and Edin, B. B. (2008). Discharges in human muscle receptor afferents during block grasping. Journal of Neuroscience, 28(48):12632– 12642. 10.1523/JNEUROSCI.3357-08.2008.

Eklund, G. (1972). Position sense and state of contraction; the effects of vibration. Journal of Neurology, Neurosurgery & Psychiatry, 35(5):606–611.10.1136/jnnp.35.5.606.

Fallon, J. B. and Macefield, V. G. (2007). Vibration sensitivity of human muscle spindles and golgi tendon organs. Muscle & Nerve: Official Journal of the American Association of Electrodiagnostic Medicine, 36(1):21–29. 10.1002/mus.20796.

Ganaie, M. A., Hu, M., Malik, A. K., Tanveer, M., and Suganthan, P. N. (2022). Ensemble deep learning: A review. Engineering Applications of Artificial Intelligence, 115:105151. 10.1016/j.engappai.2022.105151.

Geisler, W. S. and Kersten, D. (2002). Illusions, perception and bayes. Nature Neuroscience, 5(6):508–510. 10.1038/nn0602-508.

Gibson, J. J. (2014). The ecological approach to visual perception: classic edition. Psychology press.

Goodwin, G. M., McCloskey, D., and Matthews, P. (1972a). The contribution of muscle afferents to kinaesthesia shown by vibration induced illusions of movement and by the effects of paralysing joint afferents. Brain, 95(4):705–748. 10.1093/brain/95.4.705.

Goodwin, G. M., McCloskey, D. I., and Matthews, P. B. (1972b). Proprioceptive illusions induced by muscle vibration: contribution by muscle spindles to perception? Science, 175(4028):1382–1384. 10.1126/science.175.4028.1382.

Gregory, R. L. (1968). Perceptual illusions and brain models. Proceedings of the Royal Society of London. Series B. Biological Sciences, 171(1024):279– 296. 10.1098/rspb.1968.0071.

Hasan, Z. (1983). A model of spindle afferent response to muscle stretch. Journal of Neurophysiology, 49(4):989–1006. 10.1152/jn.1983.49.4.989.

Holzbaur, K. R., Murray, W. M., and Delp, S. L. (2005). A model of the upper extremity for simulating musculoskeletal surgery and analyzing neuromuscular control. Annals of Biomedical Engineering, 33:829–840. 10.1007/s10439-005-3320-7.

Houk, J. C., Rymer, W. Z., and Crago, P. E. (1981). Dependence of dynamic response of spindle receptors on muscle length and velocity. Journal of Neurophysiology, 46(1):143–166. 10.1152/jn.1981.46.1.143.

Housley, S. N., Powers, R. K., Nardelli, P., Lee, S., Blum, K., Bewick, G. S., Banks, R. W., and Cope, T. C. (2024). Biophysical model of muscle spindle encoding. Experimental Physiology, 109(1):55–65. 10.1113/EP091099.

Kanwisher, N., Khosla, M., and Dobs, K. (2023). Using artificial neural networks to ask ‘why’questions of minds and brains. Trends in Neurosciences, 46(3):240–254. 10.1016/j.tins.2022.12.008.

Kar, K. and DiCarlo, J. J. (2024). The quest for an integrated set of neural mechanisms underlying object recognition in primates. Annual Review of Vision Science, 10. 10.1146/annurev-vision-112823-030616.

Kawato, M., Hayakawa, H., and Inui, T. (1993). A forward-inverse optics model of reciprocal connections between visual cortical areas. Network: Computation in Neural Systems, 4(4):415. 10.1088/0954-898X_4_4_001.

Kissane, R. W. P., Charles, J. P., Banks, R. W., and Bates, K. T. (2023). The association between muscle architecture and muscle spindle abundance. Scientific Reports, 13(1):2830. 10.1038/s41598-023-30044-w.

Lin, C.-C. K. and Crago, P. E. (2002). Structural Model of the Muscle Spindle. Annals of Biomedical Engineering, 30(1):68–83. 10.1114/1.1433488.

Marasco, P. D. and de Nooij, J. C. (2023). Proprioception: a new era set in motion by emerging genetic and bionic strategies? Annual Review of Physiology, 85(1):1–24. 10.1146/annurev-physiol-040122-081302.

Mathis, A., Schneider, S., Lauer, J., and Mathis, M. W. (2020). A primer on motion capture with deep learning: principles, pitfalls, and perspectives. Neuron, 108(1):44–65. 10.1016/j.neuron.2020.09.017.

Mathis, M. W., Rotondo, A. P., Chang, E. F., Tolias, A. S., and Mathis, A. (2024). Decoding the brain: From neural representations to mechanistic models. Cell, 187(21):5814–5832. 10.1016/j.cell.2024.08.051.

Matthews, P. and Stein, R. (1969). The sensitivity of muscle spindle afferents to small sinusoidal changes of length. The Journal of Physiology, 200(3):723–743. 10.1113/jphysiol.1969.sp008719.

Mileusnic, M. P., Brown, I. E., Lan, N., and Loeb, G. E. (2006). Mathematical models of proprioceptors. I. Control and transduction in the muscle spindle. Journal of Neurophysiology, 96(4):1772–1788. 10.1152/jn.00868.2005.

Mumford, D. (1994). Large-scale neuronal theories of the brain, chapter Neuronal architectures for pattern-theoretic problems, pages 125–152. MIT press.

Niyo, G., Almofeez, L. I., Erwin, A., and Valero-Cuevas, F. J. (2024). A computational study of how an α-to γ-motoneurone collateral can mitigate velocity-dependent stretch reflexes during voluntary movement. Proceedings of the National Academy of Sciences, 121(34):e2321659121. 10.1073/pnas.2321659121.

Palluel, E., Aspell, J. E., and Blanke, O. (2011). Leg muscle vibration modulates bodily self-consciousness: integration of proprioceptive, visual, and tactile signals. Journal of Neurophysiology, 105(5):2239–2247. 10.1152/jn.00744.2010.

Paszke, A., Gross, S., Massa, F., Lerer, A., Bradbury, J., Chanan, G., Killeen, T., Lin, Z., Gimelshein, N., Antiga, L., et al. (2019). Pytorch: An imperative style, high-performance deep learning library. Advances in neural information processing systems, 32.

Perez Rotondo, A., Marin Vargas, A., Dimitriou, M., and Mathis, A. (2024). Modeling sensorimotor processing with physics-informed neural networks. bioRxiv, pages 2024–09.

Poppele, R. E. and Bowman, R. J. (1970). Quantitative description of linear behavior of mammalian muscle spindles. Journal of Neurophysiology, 33(1):59–72. 10.1152/jn.1970.33.1.59.

Prochazka, A. (2021). Proprioception: clinical relevance and neurophysiology. Current Opinion in Physiology, 23:100440. 10.1016/j.cophys.2021.05.003.

Prochazka, A. and Gorassini, M. (1998). Models of ensemble firing of muscle spindle afferents recorded during normal locomotion in cats. The Journal of Physiology, 507(1):277–291. 10.1111/j.1469-7793.1998.277bu.x.

Proske, U. and Gandevia, S. C. (2012). The proprioceptive senses: their roles in signaling body shape, body position and movement, and muscle force.Physiological Reviews. 10.1152/physrev.00048.2011.

Ribot-Ciscar, E., Bergenheim, M., and Roll, J.-P. (2002). The preferred sensory direction of muscle spindle primary endings influences the velocity coding of two-dimensional limb movements in humans. Experimental brain research, 145(4):429–436. 10.1007/s00221-002-1135-4.

Roll, J.-P., Albert, F., Thyrion, C., Ribot-Ciscar, E., Bergenheim, M., and Mattei, B. (2009). Inducing any virtual two-dimensional movement in humans by applying muscle tendon vibration. Journal of Neurophysiology, 101(2):816–823. 10.1152/jn.91075.2008.

Roll, J.-P. and Vedel, J. (1982). Kinaesthetic role of muscle afferents in man, studied by tendon vibration and microneurography. Experimental Brain Research, 47:177–190. 10.1007/BF00239377.

Roll, J.-P., Vedel, J., and Ribot, E. (1989). Alteration of proprioceptive messages induced by tendon vibration in man: a microneurographic study. Experimental Brain Research, 76:213–222. 10.1007/BF00253639.

Sandbrink, K. J., Mamidanna, P., Michaelis, C., Bethge, M., Mathis, M. W., and Mathis, A. (2023). Contrasting action and posture coding with hierarchical deep neural network models of proprioception. eLife, 12:e81499. 10.7554/eLife.81499.

Saul, K. R., Hu, X., Goehler, C. M., Vidt, M. E., Daly, M., Velisar, A., and Murray, W. M. (2015). Benchmarking of dynamic simulation predictions in two software platforms using an upper limb musculoskeletal model. Computer Methods in Biomechanics and Biomedical Engineering, 18(13):1445–1458. 10.1080/10255842.2014.916698.

Schaafsma, A., Otten, E., and Van Willigen, J. D. (1991). A Muscle Spindle Model for Primary Afferent Firing Based on a Simulation of Intrafusal Mechanical Events. Journal of Neurophysiology, 65(6):1297–1312. 10.1152/jn.1991.65.6.1297.

Simha, S. N. and Ting, L. H. (2024). Intrafusal Cross-Bridge Dynamics Shape History-Dependent Muscle Spindle Responses to Stretch. Experimental Physiology, 109(1):112–124. 10.1113/EP090767.

Tang, Y., Liu, J., Liu, A., Yang, B., Dai, W., Rao, Y., Lu, J., Zhou, J., and Li, X. (2023). FLAG3D: A 3D Fitness Activity Dataset with Language Instruction. In Proceedings of the IEEE/CVF Conference on Computer Vision and Pattern Recognition, pages 22106–22117.

Vargas, A. M., Bisi, A., Chiappa, A. S., Versteeg, C., Miller, L. E., and Mathis, A. (2024). Task-driven neural network models predict neural dynamics of proprioception. Cell, 187(7):1745–1761. 10.1016/j.cell.2024.02.036.

Von Helmholtz, H. (1867). Handbuch der physiologischen Optik, volume 9. Voss.

Weiss, Y., Simoncelli, E. P., and Adelson, E. H. (2002). Motion illusions as optimal percepts. Nature Neuroscience, 5(6):598–604. 10.1038/nn0602-858.

Willmore, B. and Tolhurst, D. J. (2001). Characterizing the sparseness of neural codes. Network: Computation in Neural Systems, 12(3):255. 10.1088/0954-898X/12/3/302.

Yuille, A. and Kersten, D. (2006). Vision as bayesian inference: analysis by synthesis? Trends in Cognitive Sciences, 10(7):301–308. 10.1016/j.tics.2006.05.002.

